# Mutations in Hsp40 co-chaperone change the unique canonical inter-domain interactions stimulating LGMDD1 myopathy

**DOI:** 10.1101/2025.05.02.651952

**Authors:** Ankan K. Bhadra, Geetika Aggarwal, Anshuman Jaysingh, Daniel Chen, Jil Daw, Conrad C. Weihl, Heather L. True

## Abstract

Limb-girdle muscular dystrophy D1 (LGMDD1) is a rare, dominantly inherited neuromuscular disorder caused by mutations in the HSP40 co-chaperone DNAJB6, primarily in the GF or J-domains. Currently, no treatments are available, and a challenge in understanding the disease is identifying a specific client protein for DNAJB6 in skeletal muscle. Our previous research indicated that LGMDD1 GF domain mutants in Sis1 exhibit substrate-specific effects, influenced by HSP70 activity. Herein, we found that novel mutations in the J-domain similarly affected chaperone function. The J-domain mutants exhibited variable substrate processing, reduced binding affinity to client-substrate, and decreased stimulation of Ssa1 ATP hydrolysis, with these effects being substrate-conformer-specific. Our simulation studies noted differences in inter-domain interactions linked to the mutants, which influence the Hsp40-Hsp70 ATPase cycle. These mechanistic insights enhance our understanding of LGMDD1 myopathy and help to identify potential treatment strategies in the future.

**Teaser:** Recalibrating the inter-domain interface of the mutant protein could potentially serve as a key therapeutic strategy for LGMDD1 myopathy.

## INTRODUCTION

Proteins are essential macromolecules that regulate nearly all cellular functions, requiring precise three-dimensional structures to function correctly. This complex folding process depends on a balance maintained by cellular machinery that oversees protein synthesis, folding, and degradation (*1*). Disruptions to this balance, caused by environmental or genetic stress, can lead to protein misfolding diseases (PMDs). These diseases result from the transformation of soluble proteins into insoluble amyloid deposits and include protein conformational disorders (PCDs), which encompass common neurological and muscular disorders and prion diseases (*2*). PCDs are characterized by the accumulation of amyloid aggregates, improper degradation, and amyloid polymorphism, wherein a single protein can form multiple aggregate forms that manifest different disease phenotypes, particularly in prion diseases (*3*, *4*). Cells possess defense mechanisms to combat these aggregates, with chaperone proteins playing a crucial role in disaggregating and refolding affected proteins. However, in prion diseases, chaperones may inadvertently promote the propagation of the prion state, aiding in the survival of the self-propagating structure (*5*). Mutations in these chaperone proteins can lead to various disorders known as chaperonopathies. One example of chaperonopathy is Limb-Girdle Muscular Dystrophy Type D1 (LGMDD1), caused by mutations in the DNAJB6 gene (Hsp40), a co-chaperone of Hsp70 (*6*). Specifically, LGMDD1 results from mutations in two adjacent domains of the DNAJB6 protein: the J-domain and the GF domain (*6–13*). DNAJB6 plays a vital role in protein folding and disaggregation (*14–17*); however, its specific function in maintaining skeletal muscle homeostasis is not fully understood. Moreover, the interactions between DNAJB6 and its client proteins in skeletal muscle remain unclear. The limited knowledge about the substrates of DNAJB6 poses a significant challenge in studying these myopathies. However, DNAJB clients are well-characterized in yeast, thus providing a potential model system for studying the effects of disease-causing mutants. Additionally, our understanding of DNAJB function within the yeast chaperone network is significantly more advanced than that of skeletal muscle. Previously, we employed a transdisciplinary approach to evaluate the functionality of LGMDD1-associated mutants in model systems (*18*). Using the DNAJB6 yeast homolog Sis1, we recently determined how LGMDD1 GF domain mutants alter HSP40 functions (*19*).

Hsp40s (DNAJ proteins) are ubiquitous partners for Hsp70s, collectively forming a robust chaperone system that is essential for maintaining protein homeostasis. The Hsp70 machinery interacts with client proteins through a cycle of ATP-regulated binding and release to facilitate protein folding (*20*). By determining client specificity, DNAJ proteins serve as primary facilitators of the cellular protein quality control system. They play an integral role in deciding the fate of misfolded proteins—whether they are refolded or targeted for degradation (*20*). One class of proteins that demonstrates a clear client-chaperone relationship with DNAJ family members consists of prion proteins in yeast. In contrast to pathogenic mammalian prions, yeast prions are non-toxic (*21*). Several prion proteins exist naturally within yeast, including Sup35 and Rnq1. The propagation of prions in yeast requires various protein chaperones, including Hsp104, Hsp70 (Ssa1), and the Hsp40 Sis1 (*22*). Counterintuitively, these chaperones can positively regulate prion maintenance by disaggregating prion substrates, thereby promoting seeding and transmission (*23*, *24*). Importantly, prions in yeast are not lethal, thereby allowing for phenotypic readouts in living cells that can be used to discern chaperone function. These features make yeast an ideal system to study protein aggregate formation and disaggregation as mediated by protein chaperones.

DNAJ proteins are divided into three classes based on the domains they share with the prototype *E. coli* DNAJ (*25*). In class A JDPs, the N-terminal J-domain is followed by a glycine-phenylalanine-rich region (GF), 2 homologous β-sandwich domains, βSD1 and βSD2 (previously termed CTDI and CTDII). Class A JDPs also contain a zinc-finger-like β-hairpin inserted into βSD1 and a C-terminal helical dimerization domain. Class B JDPs have a similar domain organization to Class A but generally feature a longer GF-rich region. Notably, the zinc-finger-like β-hairpin is absent in Class B. Class C JDPs only share the J-domain, which is not necessarily located at the N-terminus and can appear anywhere within the sequence (*25*, *26*). Recent data implicated the GF-rich region of DNAJB1 in an autoinhibitory mechanism that regulates the primary class B J-domain proteins (JDPs) (*27*). This inhibition can be released with second-site mutations (E50A, F102A, and or ΔH5) in DNAJB1. Our functional data suggest that the LGMDD1 GF domain mutants are not simply releasing the auto-inhibitory mechanism, however, as they show reduced binding to Hsp70 in the absence of the client, indicating that the complex (Sis1-substrate-HSP70) formed with Hsp70 is indeed driven by the Sis1-substrate interaction (*19*).

In this study, we explored the mechanistic nature of dysfunction associated with the recently identified LGMDD1 J-domain mutants in DNAJB6. We correlated these findings with those of the GF domain mutants (*19*). We generated homologous DNAJB6 LGMDD1 J-domain mutations in the essential yeast DNAJ protein Sis1 (DNAJB6-A50V (Sis1-S49V), DNAJB6-E54A (Sis1-E53A), DNAJB6-E54K (Sis1-E53K), and DNAJB6-S57L (Sis1-N56L)). We discovered that novel disease-associated variants in the Hsp40 J-domain result in abnormal chaperone function and altered protein homeostasis in a client- and conformer-specific manner in vitro. Interestingly, the J-domain mutants exhibited variability in substrate processing similar to that of the GF domain mutants. We observed a significant reduction in the binding affinity of the mutants to Hsp70 (Ssa1), both in the presence and absence of client proteins. Additionally, the stimulation of the Ssa1 ATP hydrolysis rate was significantly decreased with the mutants, although this effect was again specific to the substrate conformer. To further understand the mechanism by which J-domain mutants alter chaperone function in LGMDD1, it is essential to understand the structural interactions between chaperones and between chaperones and their client proteins. To begin to address this, we conducted *in silico* simulation studies. Our results show that when a client or Ssa1 (EEVD) binds to Sis1, inter-domain rearrangements occur within Sis1, indicating the allosteric effect of client/co-chaperone binding on all domains of Sis1. CTD-GM and J-GM interactions are lost, which are accompanied by gains in a few J-GM and J-GF interactions. J-domain mutations by themselves disrupt these unique canonical inter-domain networks within Sis1 that alter client and co-chaperone binding interfaces.

## RESULTS

### Mutants of the LGMDD1-associated homologous J-domain in Sis1 exhibit reduced efficiency in processing substrates

Our previous research delved into the roles played by LGMDD1 GF domain mutants within the chaperone protein Sis1. We employed two well-known yeast prion proteins as client substrates: Rnq1, whose Q/N-rich prion domain allows it to exist in aggregated [*RNQ+*] prion state, and Sup35, that exist as aggregated [*PSI+*] prion state (*28*). Our findings unveiled unique processing defects in the LGMDD1 GF domain mutants, which were both substrate-specific and conformer-specific (*19*). Intriguingly, these mutants supported the propagation of certain prion strains but were ineffective with others, suggesting a nuanced ability to recognize their typical clients in specific conformations but struggling to process them effectively. These findings shed new light on the complex interplay of protein misfolding and chaperone function and their implications for muscular dystrophy.

In the current study, we aimed to expand upon these findings by comparing the functional properties of LGMDD1 J-domain mutants in Sis1 with those of the wild type (WT). To achieve this, we purified the proteins (Supplementary Figure 1) and utilized them in targeted substrate processing assays conducted in vitro. Our focus was on evaluating the amyloid formation of Rnq1 in the presence of both Sis1-WT and a range of LGMDD1 J-domain mutants (Sis1-S49V, Sis1-E53K, Sis1-E53A, and Sis1-N56L). Rnq1 is recognized as a significant substrate of Sis1 and is known to form higher-order aggregates in vitro (*29*). To monitor the aggregation of Rnq1, we employed Thioflavin T (ThT), a fluorescent dye used to show the presence of the cross β-sheet conformation characteristic of amyloid fibrils (*30*). During the initial lag phase of incubation, Rnq1 monomers alone did not exhibit any significant changes in ThT fluorescence (Fig. 1A). However, we observed an abrupt increase in fluorescence, signaling the onset of aggregate formation. Notably, the presence of Sis1-WT prolonged the lag phase, indicating a stabilizing effect, whereas the LGMDD1 J-domain mutants accelerated nucleation, mirroring the behavior seen with the GF domain mutants (*19*). The time to 50% ThT fluorescence shows a strikingly similar stabilizing phenomenon of Sis1-WT compared to mutants in either the GF domain or J-domain (Fig. 1B).

**Figure 1.**
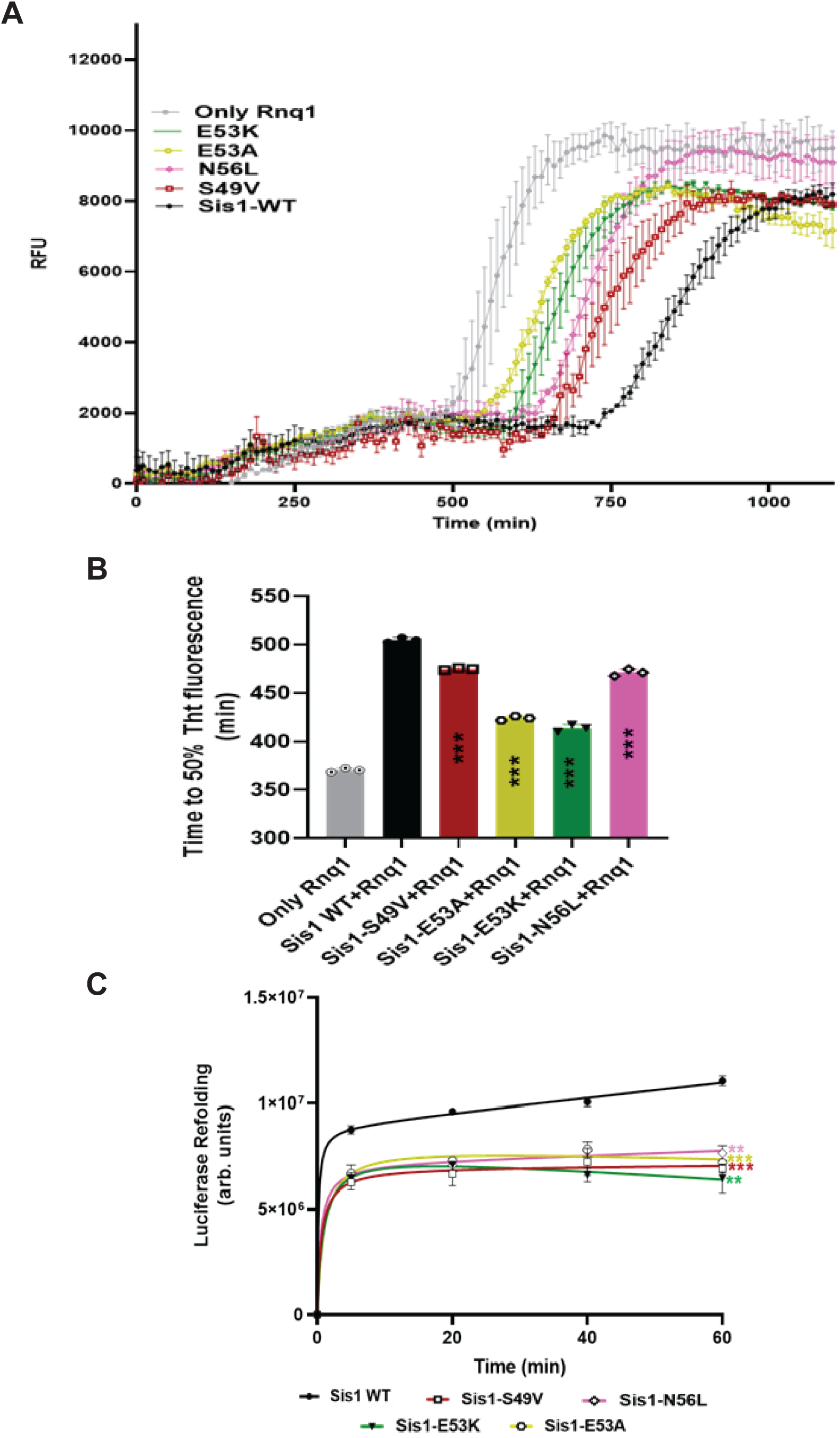
**Mutants of the LGMDD1-associated homologous J-domain in Sis1 exhibit reduced efficiency in processing substrates**. A) The kinetics of Rnq1 fibrillation in the presence of unseeded Rnq1 only (grey), Sis1-WT (black), Sis1-S49V (red), Sis1-E53A (yellow), Sis1-E53K (dark green), or Sis1-N56L (dark pink) were measured using a ThT fluorescence assay. (B) The time to 50% ThT fluorescence was calculated by fitting the graph using the EC50 equation (Y = Bottom + (Top-Bottom)/(1 + 10^ ((LogEC50-X) *Hillslope) in GraphPad Prism. (C) Refolding activity of heat-denatured luciferase in the presence of LGMDD1 mutants. Luciferase and Ssa1-WT and Sis1-WT/mutants were incubated at 42 °C for 10 min to heat denature luciferase. At various time points, activity was measured by a luminometer after adding substrate. The mean ± SEM values, n = 3 biologically independent samples are shown. For (C), each LGMDD1 mutant was compared with Sis1-WT across all time points, and *** p < 0.001 and **p < 0.01 values are reported for an unpaired, two-sided t-test. Data represented as mean values ± SEM, n = 3 biologically independent samples.

Next, we aimed to investigate whether the ability of mutant chaperones in processing substrates could be enhanced with the assistance of co-chaperones. Previous research has shown that Sis1, when lacking the GF domain (Sis1ΔGF) or possessing mutations in the LGMDD1 homologous GF domain, demonstrated a significant reduction in chaperone activity (*19*, *31*), emphasizing the importance of the GF domain for chaperone function (*19*, *31*). Herein, we evaluated the capacity of LGMDD1 J-domain mutants to assist in client refolding by conducting a luciferase refolding assay. Specifically, we heat-denatured luciferase and observed its refolding in the presence of chaperones. The results indicated that all LGMDD1 J-domain mutants were impaired in their ability to refold luciferase (Fig. 1C).

### Mutants of the LGMDD1-associated homologous J-domain in Sis1 alter substrate and Hsp70 binding

Client processing by Hsp40 is driven by two key factors: the initial binding of Hsp40 to the client substrate and its subsequent interaction with the Hsp70 co-chaperone. To investigate these interactions, we used two different substrates: full-length Rnq1 (FL Rnq1), and the enzyme luciferase. We conducted an Enzyme-linked Immunosorbent assay (ELISA) to evaluate the binding dynamics in a manner that was previously successful in the examination of the interactions between the E. coli DnaJ protein and its substrates (*32*). Our findings indicate that the mutants of the LGMDD1 J-domain exhibit significantly different binding efficiencies when interacting with denatured FL Rnq1, and luciferase, particularly in comparison to the Sis1-WT (Fig. 2A, and B). The second critical factor in Hsp40’s client processing efficacy is its interaction with the Hsp70 co-chaperone. Previous work demonstrated that the detrimental effects of DNAJB6 GF domain mutants are Hsp70-dependent (*33*) and that LGMDD1 GF domain mutants impacted the productive association with Hsp70 (*33*). Additionally, reducing the interaction between LGMDD1 mutants and Hsp70 using pharmacologic compounds led to muscle strength and myopathology improvements in mouse models (*33*). The J-domain mutants may influence their association with Hsp70 by either reducing binding affinity, sequestering Hsp70, or being hyperactive, thus altering the refolding process. Binding assays were performed to determine the association between LGMDD1 mutants and Ssa1, revealing a significant reduction in interaction with Ssa1-WT both in the absence (Fig. 2C) and presence of client protein Rnq1 (Fig. 2D). This indicates that the interaction defect between the mutants and Ssa1-WT is client-independent, as was observed with GF domain mutants (*19*).

**Figure 2.**
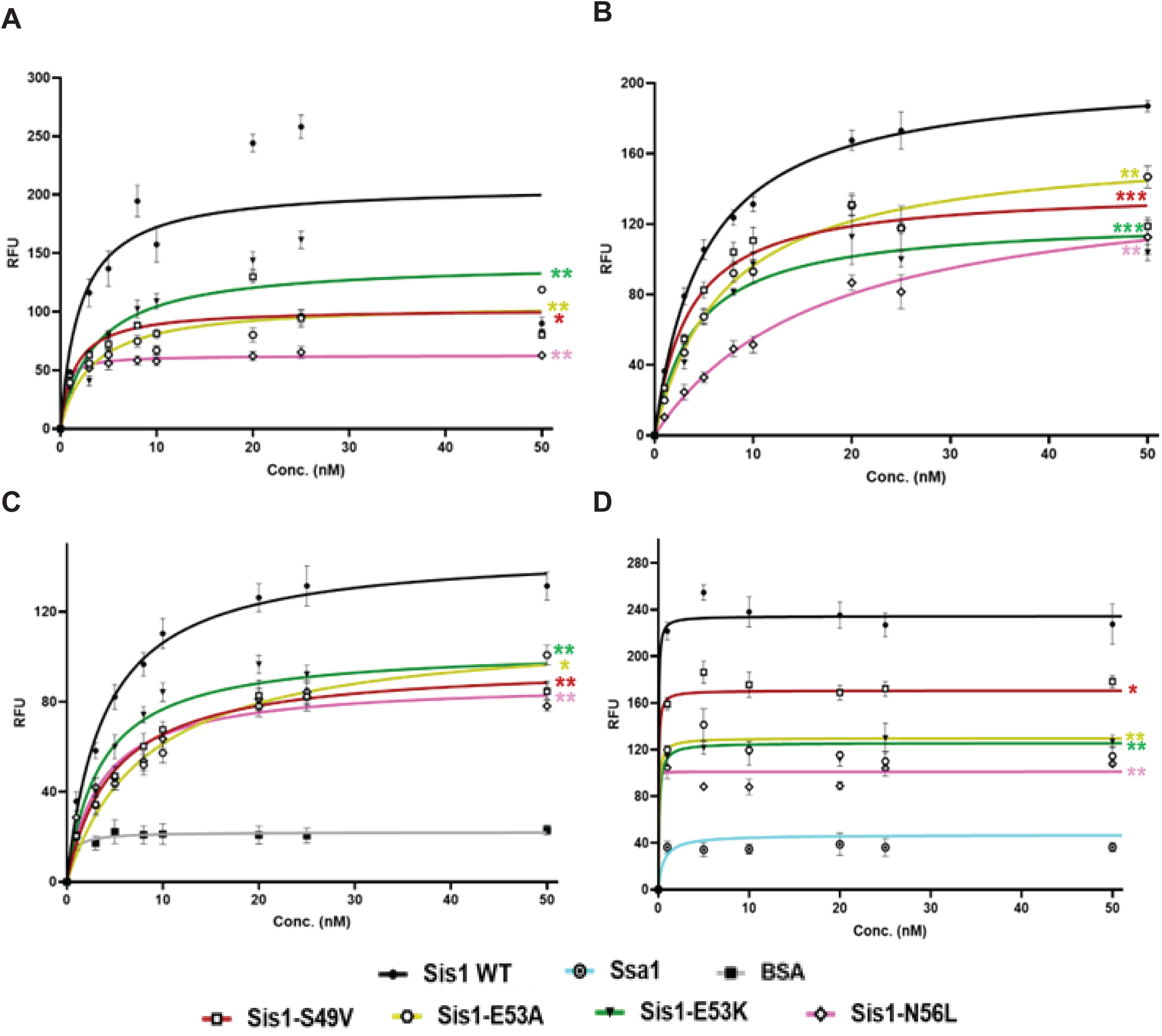
**Mutants of the LGMDD1-associated homologous J-domain in Sis1 alter substrate and Hsp70 binding**. Binding of purified Sis1-WT (black), Sis1-S49V (red), Sis1-E53A (yellow), Sis1-E53K (dark green), or Sis1-N56L (dark pink) to denatured full-length (FL) Rnq1 (A), and luciferase (B). FL Rnq1 (400 ng) and luciferase (100 ng) were immobilized in microtiter plate wells, and dilutions of purified Sis1-WT and Sis1-mutants (0, 1, 3, 5, 8, 10, 20, 25, 50 nM) were incubated with each client. The amount of Sis1 retained in the wells after extensive washing was detected using a Sis1-specific antibody. For (C); Ssa1 (200 nM) was immobilized in microtiter plate wells and dilutions (0, 1, 3, 5, 8, 10, 20, 25, 50 nM) of purified Sis1-WT (black), Sis1-S49V (red), Sis1-E53A (yellow), Sis1-E53K (dark green), Sis1-N56L (dark pink), or BSA (grey) as a control, were incubated with it. Bound Sis1 was detected using an αSis1 antibody. Denatured Rnq1 (D) was premixed with Sis1-WT/mutants and immobilized in microtiter plate wells. Dilutions of Ssa1-WT (0-50 nM) were then incubated with it. Bound Ssa1-WT was detected using an αSsa1 antibody. Only Ssa1 (cyan blue) was used as a control for (D). Data represented as mean values ± SEM, n = 3 biologically independent samples.

### Mutants of the LGMDD1-associated homologous J-domain in Sis1 alter its ability to function efficiently with Ssa1 (Hsp70)

Heat shock proteins (Hsp40s and Hsp70s) work together as co-chaperones in cellular processes. Hsp40s play a crucial role in enhancing the ATPase activity of Hsp70s, which, in turn, significantly influences how client substrates are handled within the cell [20]. The ability of these chaperones to recognize client substrates is largely determined by the conformational state of the substrates. Previous studies have suggested that the GF domain of Hsp40s is instrumental in this conformation-dependent recognition (*34–37*). In our earlier research, we identified specific functional defects associated with client conformers that are linked to mutations in the GF domain of the LGMDD1 proteins, observed in both in vivo and in vitro studies (*18*, *19*). As such, we aimed to explore how the conformation of clients influences the functionality of these mutants. To investigate this, we measured the capacity of the Sis1-WT and various LGMDD1 J-domain mutant proteins to stimulate the ATPase activity of Ssa1, both with and without the presence of different conformers of the Rnq1 protein. We standardized the concentration of Rnq1 monomers, as described previously (*19*), and conducted a phosphate standard curve for each assay (Supplementary Figure 2A). Interestingly, our results revealed no significant differences in the stimulation of Ssa1 ATP hydrolysis between Sis1-WT and the LGMDD1 J-domain mutants when client proteins were absent (Supplementary Figure 2B) or present as Rnq1 monomers (Fig. 3A). Similarly, there was no variation in ATPase activity when Rnq1 amyloid formed at 37 °C were included (Fig. 3B). However, in the presence of Rnq1 amyloids formed at lower temperatures, specifically 18 °C and 25 °C, we observed a marked reduction in stimulation of Ssa1 ATPase activity by the LGMDD1 mutants (Fig. 3C-D). This suggests that “denatured monomers” provide a diverse array of epitopes that chaperones can effectively recognize. By contrast, amyloid structures likely present a more limited number of chaperone recognition sites. These findings indicate that the different conformations of Rnq1 can create distinct interactions with Sis1, leading to varying levels of Hsp70 ATPase stimulation. Furthermore, the LGMDD1 mutants consistently demonstrated functional deficiencies, suggesting that alterations in client conformation can significantly impact chaperone activity.

**Figure 3.**
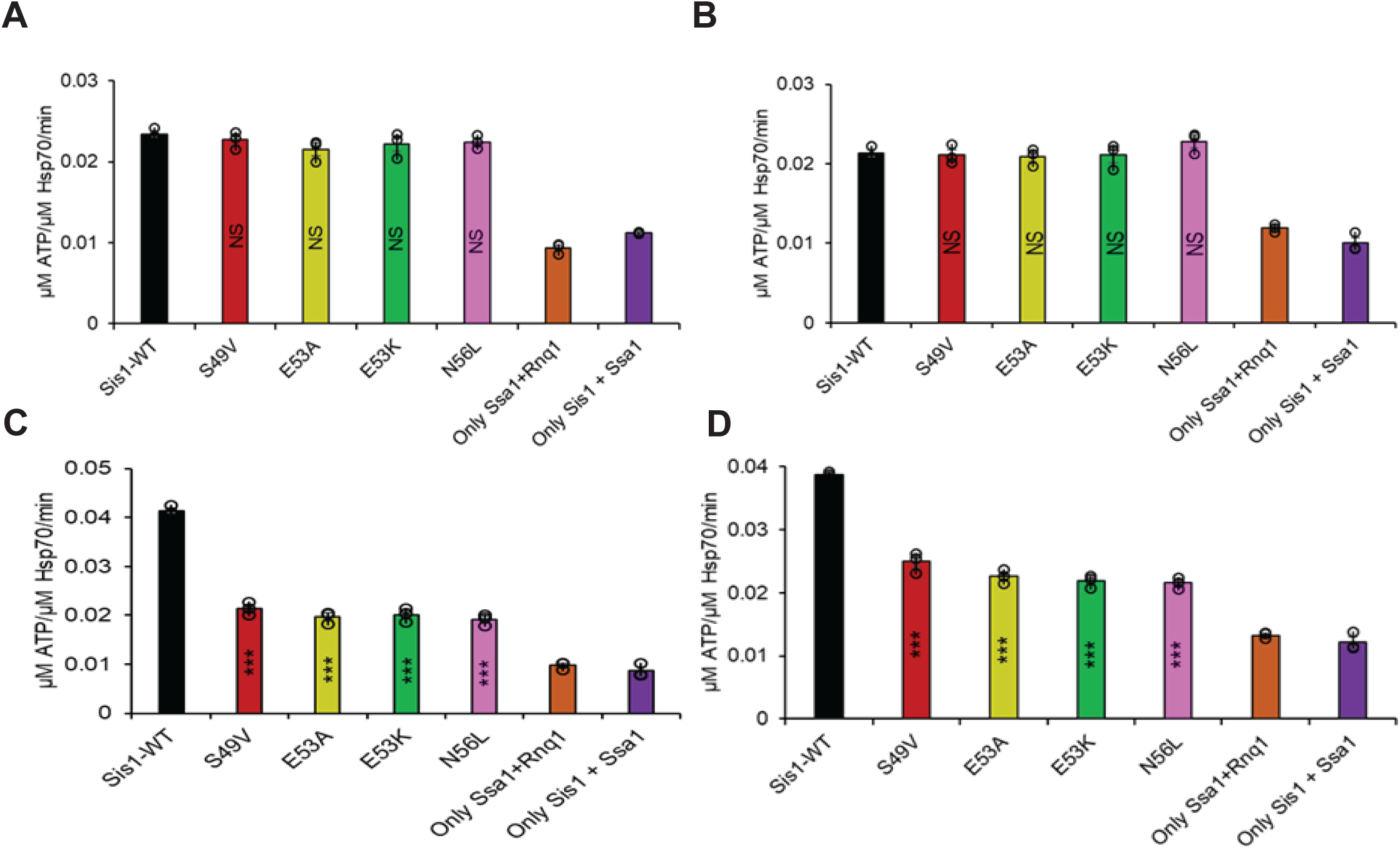
**Mutants of the LGMDD1-associated homologous J-domain in Sis1 alter its ability to function efficiently with Ssa1 (Hsp70)**. Stimulation of Ssa1 ATPase activity in the presence of Rnq1 monomer as client (A), and Rnq1 seeds formed at (B) 18°C, (C) 25°C, and (D) 37°C as clients. Ssa1 (1 μM) combined with ATP (1 mM) in the presence of Sis1-WT (black) or Sis1-mutants (Sis1-S49V, red; Sis1-E53A, yellow; Sis1-E53K, dark green; or Sis1-N56L, dark pink, 0.05 μM). The fraction of ATP converted to ADP was determined. For (A), the Rnq1 monomer used was 25 μM. For (B), (C), and (D), a total of 10% of the seeds were used in a reaction. Only Ssa1 with Rnq1 (orange) and Sis1 with Ssa1 (dark purple) were used as controls for A-D. Data represented as mean values ± SEM, n = 3 biologically independent samples.

### Homology Modeling reveals a compact full-length Sis1 structure consistent with NMR assignments

A full-length crystal structure of Sis1 remains unresolved, owing to the intrinsic flexibility of the glycine-phenylalanine (GF) and glycine-methionine (GM) rich domains. To enable mechanistic insights into Sis1’s interactions with client proteins and the Hsp70 homolog Ssa1, we generated full-length homology models of Sis1. Homology models were built using three approaches: AlphaFold3, Robetta, and SWISS-MODEL (*38–40*). AlphaFold3 and Robetta each generated five models, while SWISS-MODEL produced a single model based on the highest sequence identity templates available. All models were structurally aligned to the available crystal structures of the isolated J-domain and peptide-binding domains of Sis1. Structural comparisons were evaluated based on backbone root-mean-square deviation (RMSD) and domain organization relative to the crystal structures (Supplementary Table 1). The model derived from AlphaFold3 exhibited the highest structural congruence to experimentally determined domains and was selected for further analysis (Supplementary Figure 3). Notably, NMR resonance assignments covering ∼75% of Sis1’s backbone, including the disordered GF and GM regions, have recently become available (*41*). These studies identified a stable α-helix spanning residues 107-119 within the otherwise flexible GF region, consistent with the secondary structure predictions in our selected model. Site-directed mutants S49V, N56L, E53A, and E53K, all localized to Helix II of the J-domain and located near the highly conserved HPD motif (Supplementary Figures 3B and C), were computationally generated using CHARMM-GUI (*42*, *43*). Following model building, all wild-type and mutant structures were subjected to energy minimization and equilibration using GROMACS (version 2020.4) (*44*) on the Bridges2 HPC cluster. These equilibrated structures were the starting point for 1 μs all-atom molecular dynamics (MD) simulations.

### J-Domain mutations change inter-domain networks and the compactness of Sis1

Consistent with previous findings for the Hsp40 family member DNAJB6b (*45*), analysis of the radius of gyration (Rg) revealed dynamic fluctuations of Sis1 within the 25–35 Å range (Supplementary Figure 4A). The S49V and N56L mutants maintained Rg distributions similar to wild-type Sis1 (Sis1WT), while S49V exhibited a noticeable reduction in Rg fluctuations, suggesting increased molecular compactness attributable to the hydrophobic substitution at position 49. By contrast, the E53A and E53K mutants displayed increased Rg values with more frequent and pronounced excursions, indicative of an expansion in the overall molecular dimensions and a destabilization of the compact conformation (Supplementary Figure 4A). Rg histograms revealed that S49V and N56L mutants sampled a conformational ensemble similar to Sis1WT, whereas E53A and E53K mutants adopted a broader distribution of conformations (Supplementary Figure 4B).

Comparative contact map analyses (Supplementary Figure 4 and Figure 5) further revealed that while the intradomain secondary structure of the J-domain and CTD domains was preserved across all variants, substantial perturbations were observed in the inter-domain interaction profiles. In Sis1WT, Helices II and III of the J-domain extensively contacted the GF domain and portions of the GM domain (Supplementary Figure 5B). Recent MD studies on DNAJB6b indicate that the closed conformation is predominant during the simulation (*45*). The J-GF interactions observed in Sis1 resemble those found in the closed conformation of DNAJB6b (*45*). These specific contacts were largely abolished across all mutants (Figure 4A-D), particularly the interaction between Helix III and F100 of the GF domain (Figure 4E-H, green arrows). Moreover, Sis1WT exhibited stable interactions between the J-domain and the CTDI domain (Supplementary Figure 5D), and these interactions were entirely lost in the mutations (Figure 4M-P). Compensatory increases in GF– CTD and GM–CTD interactions were observed, particularly in the S49V and N56L mutants (Figure 4I-L), though they were only partial in E53A and E53K. Notably, the interactions between the HPD motif and the GF domain were also somewhat perturbed (Supplementary Figure 6A-D).

**Figure 4.**
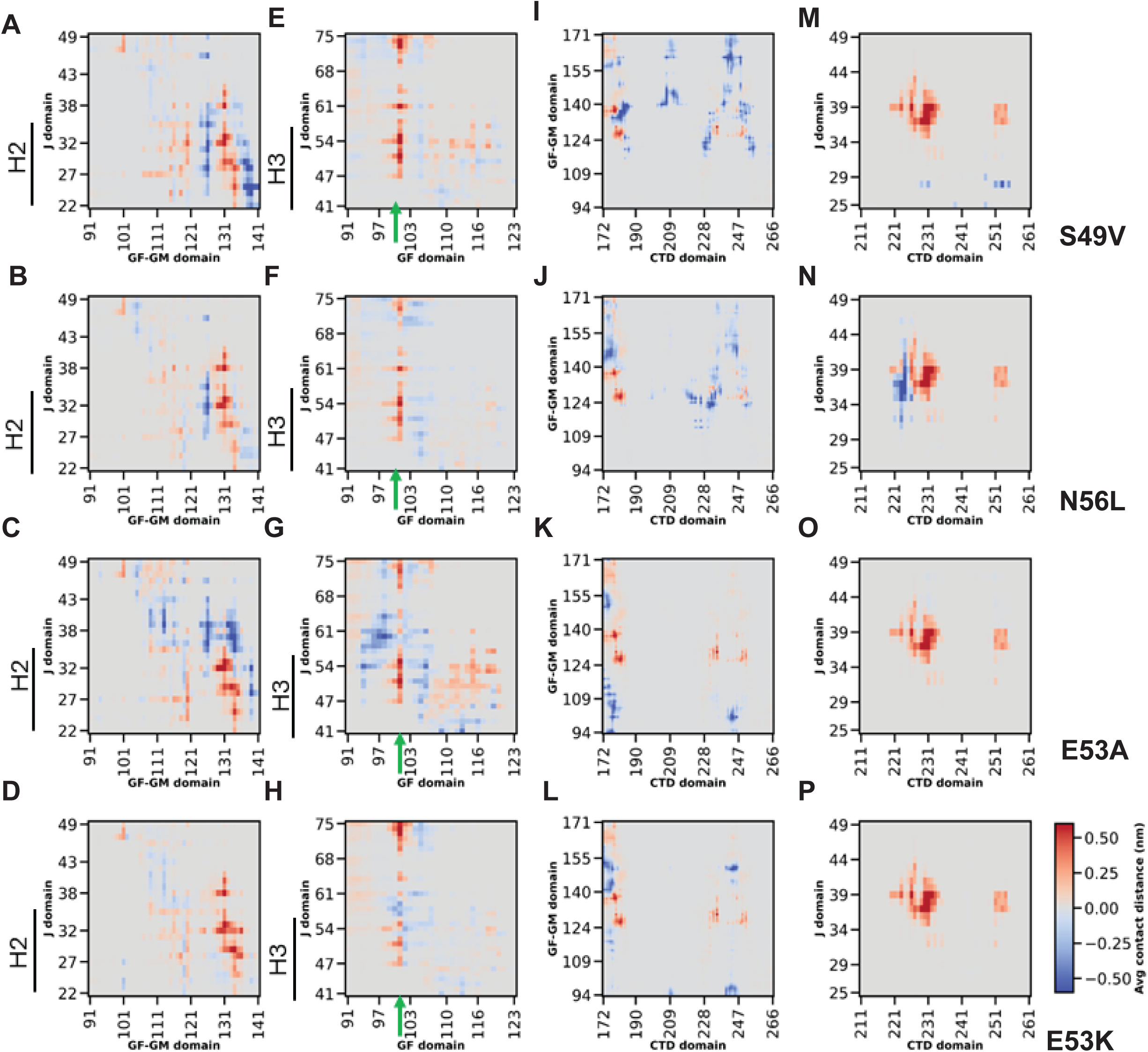
**J-domain mutations selectively disrupt Sis1 interdomain connectivity**. Changes in the interdomain contacts of Sis1 mutants are shown after subtracting WT contacts to generate difference-contact maps (plotted as average contact distance for different domains). Positive values reflect contact loss (red); negative values indicate the formation of new interactions (blue). (A-D) J (H2 helix)–GM contacts (notably residues 130–138) are reduced in all mutants, weakening interdomain coupling. (E-H) Reduced persistence of J–GF contacts (notably around residues 100–101 in the GF region) indicates compromised communication between the H3 helix of J and GF domains. (I-L) S49V and N56L mutations enhance GM–CTD interactions. However, this enhancement is minimal with E53 mutations. (M-P) J–CTD long-range contacts observed in WT are universally lost in all mutants, suggesting a common destabilization pathway across variants. Contact maps computed using Conan 1.1.

**Figure 5.**
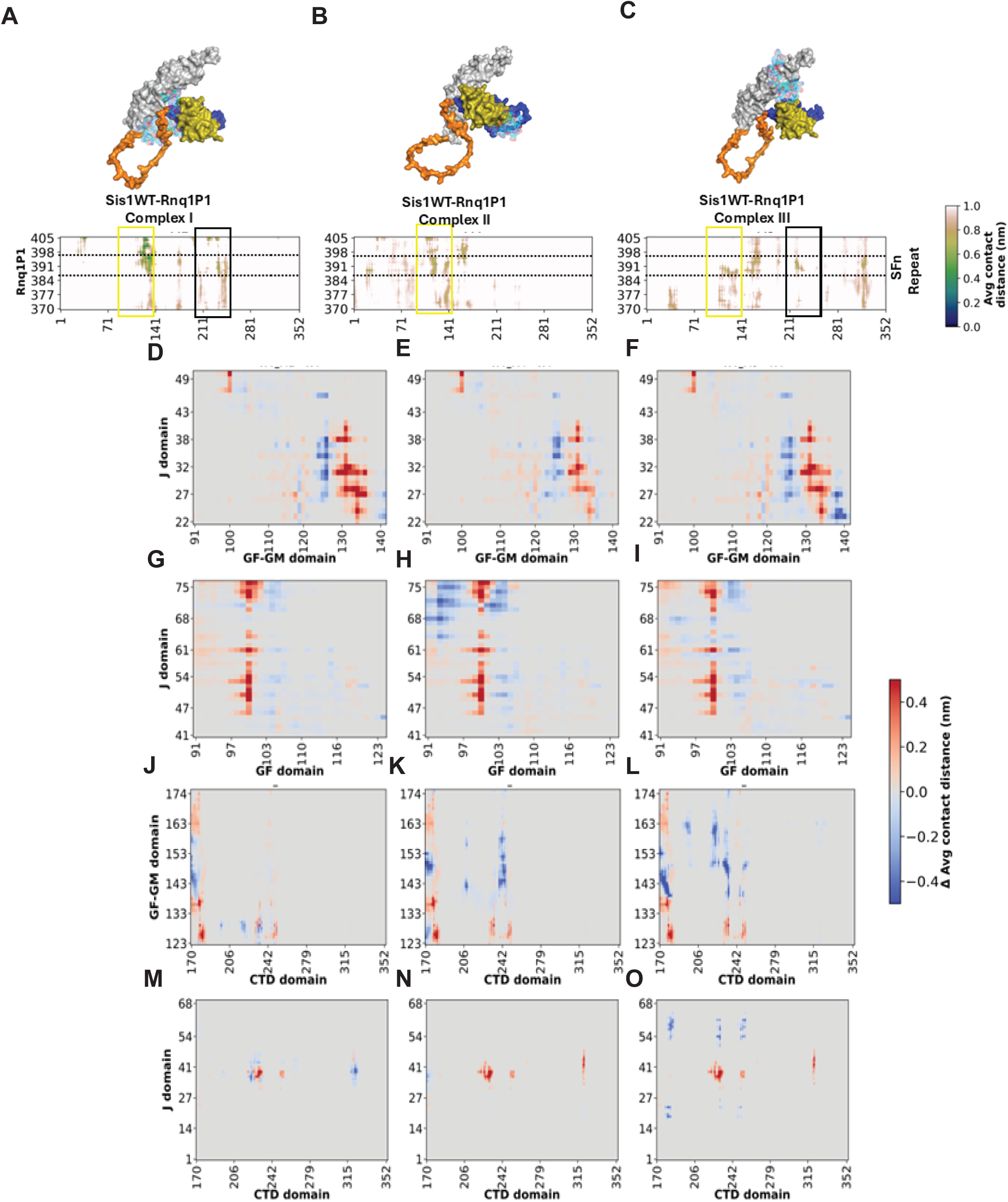
**Rnq1 Binding to Sis1 drives inter-domain interactions similar to J-domain mutations**. Sis1WT is rendered in surface representation; domains are color-coded (J domain (olive), GF (dark blue), GM (orange), and CTD domain (grey) are shown). Peptide Rnq1P1 is displayed as a cyan mesh. (A-C) Contact maps of Sis1WT–Rnq1P1 across three top-ranked binding poses. (D-O) Inter-domain contacts of Sis1WT-Rnq1P1 complexes are subtracted in Sis1WT only across three docking modes, and difference-contact maps were plotted as average contact distance for different domains. Positive values reflect contact loss (red); negative values indicate the formation of new interactions (blue). Sis1 binding to client peptide disrupts inter-domain interactions similar to those between J-GF, J-GM, and GM-CTD, while J-CTD contacts are identical to those of mutants in the absence of a client.

Collectively, our results indicate that the interaction between different domains is primarily lost due to the LGMDD1 J-domain mutations. Our next question to address was whether these losses of inter-domain interactions are responsible for the reduced binding between Sis1 and the client and/or Sis1 and the EEVD motif of Hsp70. A recent study on DNAJB8 showed that the inter-domain interactions drive Hsp70 recruitment to DNAJB8 (*46*), highlighting the importance of inter-domain interactions in the Hsp40-Hsp70 chaperone refolding system. Moreover, NMR studies on other DNAJB proteins and Hsp70 suggest that changes in J-GF domain interactions allow the substrate binding domain (SBD) of Hsp70 to bind to the GF domain (*47*). Our results also show that even distant GM-CTD interactions are changed due to J-domain mutations, indicating the involvement of long-range allosteric effects. As such, we hypothesize that the J-domain mutations reduce the client and EEVD binding to Sis1 due to the changes in inter-domain interactions.

### J-Domain mutations reconfigure Rnq1 binding hotspots on Sis1

Rnq1, a canonical substrate of Sis1, is characterized by a predominantly disordered C-terminal region (amino acids 154–405) containing tandem QN repeat motifs along with multiple SALASMASSY repeats and a unique SFn motif (SFNFSGGNFS) located within the major structure of the Rnq1 prions, C-terminal Core 1 (*48*). This region is unique because the mutation of this region impairs the propagation of all [*RNQ+*] variants (*49*). To specifically probe the SFn repeat-mediated interaction, we selected a 36-residue peptide from Rnq1 (residues 370–405), hereafter termed Rnq1P1. MD simulations were conducted on the AlphaFold3-predicted structure of FL Rnq1 (PDB ID: AF-P25367-F1-model_v4, Supplementary Figure 7A) to generate an ensemble of RNQ1p1 conformations. Structural clustering based on secondary structure similarity identified ten representative conformations (Supplementary Figure 7B inset). These conformations were docked onto Sis1 using HDOCK (*50*), and the resulting complexes were analyzed by contact map analysis. Rnq1p1 exhibited widespread binding across the Sis1 surface, with dominant interaction hotspots around the GF domain and secondary binding to the CTD (Supplementary Figure 7C-D). This binding pattern is consistent with recent NMR studies of Sis1-client interactions (*51*). Finally, 500 ns MD simulations were performed on selected docked complexes where Rnq1p1 engaged the GF and CTD domains. Corresponding S49V and E53A mutations were introduced into these complexes using CHARMM-GUI, followed by energy minimization and equilibration.

### Rnq1 binding to Sis1 drives inter-domain interactions similar to J-domain mutation

MD trajectory analysis of the top 3 Sis1WT-Rnq1P1 complexes showed that throughout the MD run, Rnq1P1 remains bound near the CTD and GF domains in all complexes (Figure 5A-C). Their corresponding contact maps showed that in complexes I and III, the peptide binds to CTD (black boxes) and near the GF domain (yellow boxes). The peptide predominantly binds in complex II at the GF domain (yellow box). These regions span from 119-140 in the GF region and 225-240 in the CTD region. Upon analyzing the inter-domain interactions when Rnq11P1 peptide binds to Sis1 in each complex (Figure 5D-O), we found that the H2 and H3 helices of the J-domain, which interact with the GF and GM domains, are lost when Rnq1P1 peptide binds to Sis1 (Figure 5D-I). Interestingly, this loss is similar to that observed when the J-domain mutation is introduced in Sis1 (Compare Figure 5D-I and 4A-H). Likewise, the GF/GM-CTD and J-CTD interactions are similarly lost in the mutants alone and when the Rnq1 peptide binds to Sis1 (Compare Figure 5J-O and Figure 4I-P). This indicates that when Rnq1 binds to Sis1 in normal conditions, these inter-domain interactions generally occur, suggesting a specific and unique allosteric pathway for client binding, which is disrupted by the LGMDD1 J-domain mutations.

Furthermore, J-domain mutation reduces the interaction between Rnq1 peptide and Sis1 (Figure 6A-F; green boxes); however, additional interactions were seen between peptide and mutants (Figure 6A-F; green boxes). We performed a binding assay to investigate these interactions using Rnq1 lacking the N-terminal region (ΔN Rnq1) as substrate. We found that the mutants of the LGMDD1 J-domain exhibit a minimal yet significant reduction in binding to ΔN Rnq1 as the substrate, as compared to the Sis1-WT (Figure 6G). This finding agrees with our structural simulation data (Figure 6 A-F). This (Figure 6G), along with our binding assay data (Figure 2A, B), reinforces the notion that these J-domain mutants operate in a substrate-specific manner, suggesting that their effectiveness varies depending on the specific substrate being processed.

**Figure 6.**
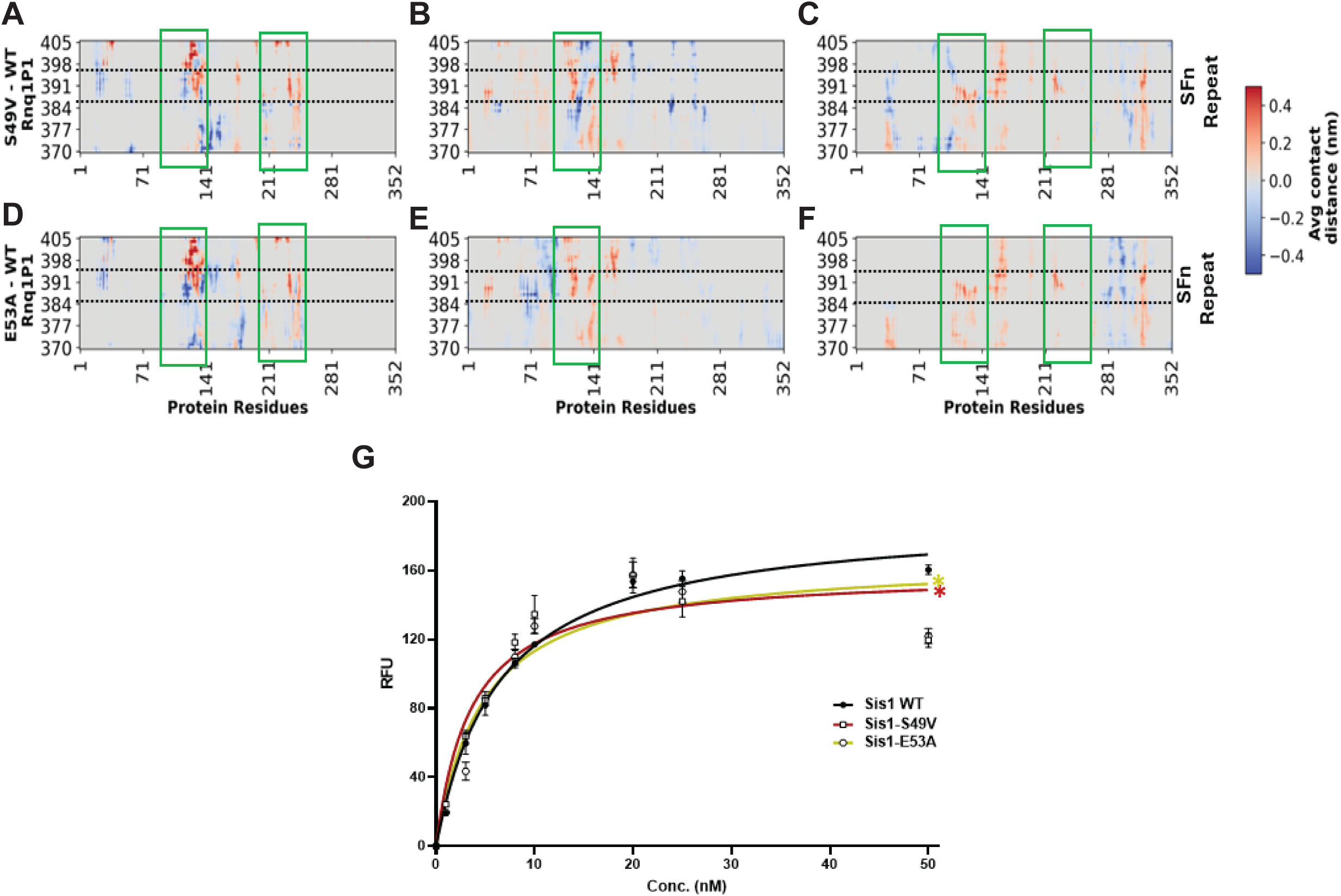
**S49V and N56L mutations alter the binding interface between Sis1 and Rnq1P1 across three docking modes**. Difference-contact maps (plotted as average contact distance) were generated by substracting Sis1 WT-Rnq1P1 from mutant-Rnq1P1 complexes. Positive values reflect contact loss (red); negative values indicate the formation of new interactions (blue), are shown in green boxes. (A-C) Difference-contact maps for S49V–Rnq1P1 complexes reveal weakened contacts across the J and GF domains. (D-F) Difference-contact maps for N56L–Rnq1P1 complexes show altered interaction topography, with increased contacts near the GM domain and decreased contacts in the J-domain region. (G) Binding of purified Sis1-WT (black), Sis1-S49V (red), Sis1-E53A (yellow) to denatured ΔN Rnq1, which contains Rnq1P1. ΔN Rnq1 (400 ng) was immobilized in microtiter plate wells, and dilutions of purified Sis1-WT and Sis1-mutants (0, 1, 3, 5, 8, 10, 20, 25, 50 nM) were incubated with each client. The amount of Sis1 retained in the wells after extensive washing was detected using a Sis1-specific antibody. For G, each LGMDD1 mutant was compared with Sis1-WT across all time points, and *p < 0.05 values are reported for an unpaired, two-sided t-test. Data represented as mean values ± SEM, n = 3 biologically independent samples.

### J-Domain mutations impair multisite Ssa1 (EEVD) binding on Sis1

J-domain proteins like Sis1 enable Hsp70s like Ssa1 to refold newly synthesized denatured or aggregated proteins (*52*). Sis1 binds to clients and delivers them to Ssa1 (*52*). Ssa1 can bind to Sis1 alone; however, these interactions are very dynamic [46]. Sha et al 2006 showed that the EEVD motif present at the C-terminal of Ssa1 binds to the CTD domain of Sis1, the only part of a complex that was stably captured in the crystal structure of the Ssa1-Sis1 complex (*53*). To investigate the effect of J-domain mutations on the binding of Ssa1’s EEVD motif to Sis1, we employed AlphaFold3-based peptide-protein docking. In all of the top-ranked docked complexes, the EEVD peptide engaged the CTDI domain, a finding that is in agreement with previous crystallographic and NMR studies (*51*, *53*). The 1 μs long equilibrium MD simulations confirmed the stability of Ssa1 (EEVD) binding in the WT complex, predominantly through residues spanning 180-186, 197-204, and 230–257 (Figure 7A, marked three arrows). The J-domain mutations selectively disrupted interactions at the 180-186 and 230-257 segments, while maintaining contacts at the middle 197-204 region (Figure 7B-C). Kullback-Leibler (KL) Divergence analysis (*54–56*) was performed to further analyze protein-peptide interactions in wild type and J-domain mutants. KL Divergence represents regions of protein-peptide in the mutant where the mutation has caused significant changes in the interaction patterns. KL divergence analysis between WT and mutant contact maps highlighted a marked redistribution of interaction probabilities, particularly at residues 180-186 (Figure 7B). Differential contact maps corroborated these findings, identifying a net loss of EEVD-CTD contacts in the mutants (Figure 7C).

**Figure 7.**
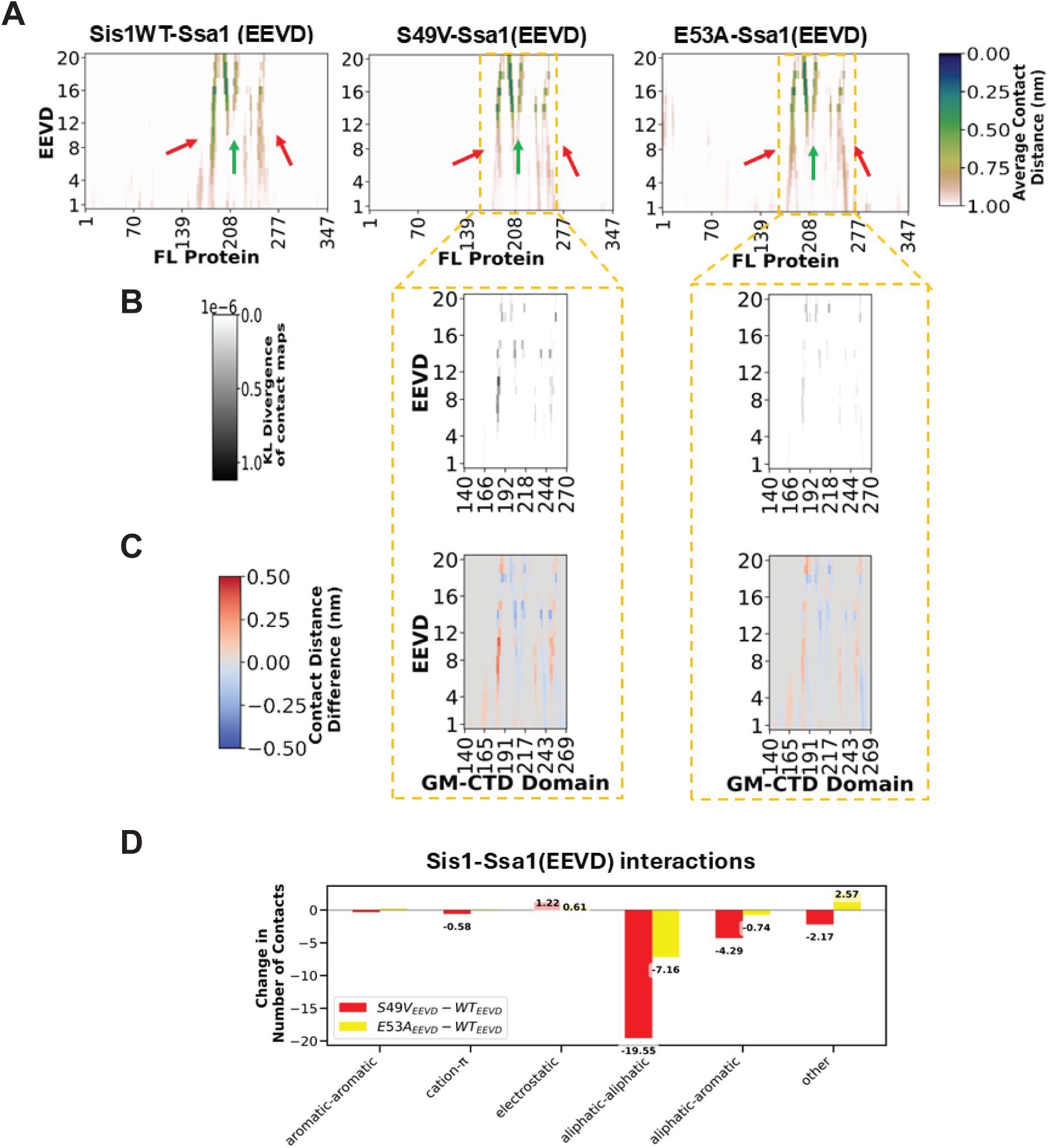
**Hsp70 EEVD motif interaction sites on Sis1WT and mutants involve the CTD but display mutation-specific dynamic reorganization**. (A) Contact maps reveal EEVD binds predominantly at CTD residues, with interaction persistence visualized via grayscale intensity. The average contact distance was computed, and the dark color represents high contact persistence, and the light color represents low contact persistence. EEVD binds to Sis1WT and mutants through three regions on the CTD (marked by green and red arrows) (B) Kullback-Leibler (KL) divergence of contact frequencies identifies mutation-sensitive residues exhibiting enhanced sidechain mobility during EEVD engagement. (C) Difference-contact maps highlight mutation-specific loss or gain in EEVD-binding residues. (D) The average number of Sis1-Ssa1 (EEVD) contacts was calculated between Sis1WT/mutants and EEVD, categorized by residue types. Aliphatic-aliphatic and aliphatic-aromatic interactions dominate binding. Difference in the average number of contacts revealed that both S49V and E53A display a net reduction in hydrophobic interactions.

To understand how these residues are affected in mutants, we computed the average contact profile of these residues in Sis1WT and mutants on binding to EEVD (Supplementary Figures 8 and 9). In Sis1WT-Ssa1 (EEVD), 180-186 residues generally interact with the CTD domain and Ssa1 (EEVD) (Supplementary Figure 8A). The average difference contact profile was calculated by subtracting the average contacts of WT from each mutant. Subsequent analysis revealed that, in mutants, residues 180-186 established non-specific intramolecular interactions with GF and GM domains (Supplementary Figure 8B), thereby decreasing their availability for EEVD engagement.

Next, we checked whether the J-domain mutation affects HPD intermolecular interactions with the GF domain within Sis1 upon binding to the EEVD motif, because previous reports showed that EEVD binding releases GF autoinhibition, which allows the HPD motif to interact with Ssa1 and subsequent ATP hydrolysis (*27*, *45*, *57*). Our results show that a J mutation doesn’t affect HPD motif interdomain interactions with the GF domain (Supplementary Figure 10A-B). These observations are supported by our ATP hydrolysis activity of Ssa1, which remains the same in the presence of Sis1 WT and its mutants without any client (Supplementary Figure 2B). This indicates that reduced binding between Ssa1 and J-domain mutants might be partly due to reduced EEVD-CTD interactions.

### Disruption of interdomain contacts when J-domain mutants bind to client and Ssa1-EEVD

When Sis1WT binds to Rnq1P1, interdomain changes occur in Sis1 (Supplementary Figure 11). Specific GM residues (spanning 125-139) that bind to the CTD domain and H1 helix of J-domain in Sis1 are lost on binding to Rnq1P1 (Supplementary Figures 11A-C and 11D-F, respectively). The free GM domain in the Sis1-Rnq1P1 complex forms new interactions with the G-rich region of the GF domain (Supplementary Figures 11G-I). We observed that the G-rich region also binds to the H4 helix of the J-domain in the complex (Supplementary Figure 11J-L). A similar interaction pattern was observed when Ssa1 (the EEVD motif) engages with Sis1 (Supplementary Figure 12). This is in accordance with a recent study where the EEVD motif compete with the client binding site on Sis1 (*51*). The binding of either client or EEVD was expected to drive similar changes in the interaction pattern in Sis1 because both have a binding site near or on the CTD domain. However, our difference-contact map analysis showed that changes in inter-domain contacts in mutant-Rnq1P1 and mutant-Ssa1(EEVD) are mutation-specific (Figure 8). For instance, in the S49V mutation, there is a strong non-regulated loss of interaction between GM-CTD and GM-J-domain (Figure 8A, B, E, F), which is minimal with the E53A mutation (Figure 8I, M, J, N). Interestingly, the interactions of the GM and J-domain with the G-rich region of the GF domain are lost in both Sis1-Rnq1P1 and Sis1-Ssa1 (EEVD) complexes, irrespective of the type of mutation (Compare Figure 8C, D, G, H with Figure 8K, L, O, P).

**Figure 8.**
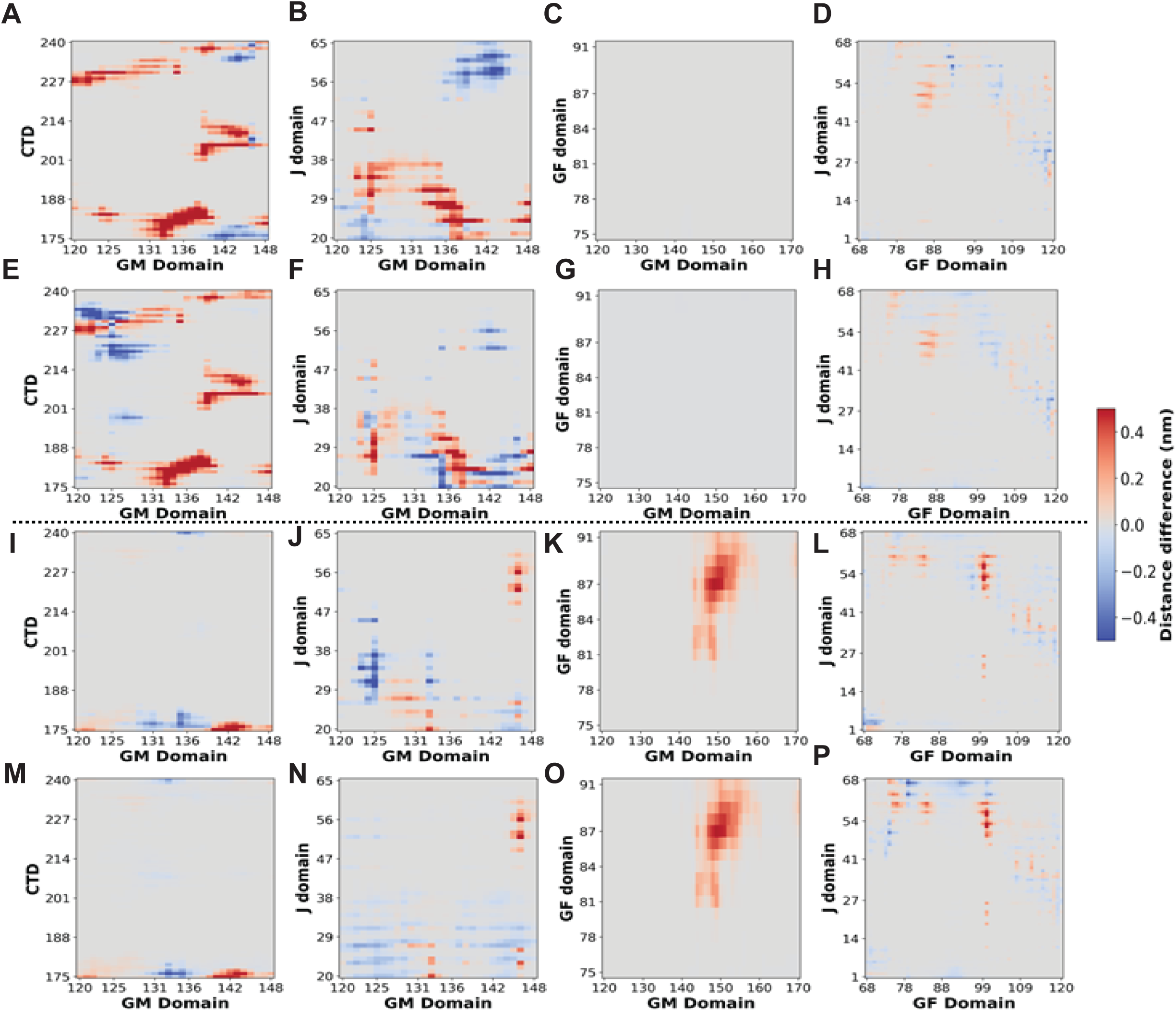
Disruption of interdomain contacts when the client and Ssa1 (EEVD) bind to mutants. The upper panel represents difference-contact maps which are generated by subtracting S49V mutant’ contacts from S49V in complex either with client (A-D) or Ssa1 (EEVD) (E-H). The bottom panel represents difference-contact maps which are generated by subtracting E53A mutant’ contacts from E53A in complex with either client (I-L) or Ssa1 (EEVD) (M-P). Difference-contact maps were plotted as the average contact distance for different domains. Positive values reflect contact loss (red); negative values indicate the formation of new interactions (blue).

### Proposed role of inter-domain interactions in the Hsp40-Hsp70 functional cycle with respect to LGMDD1 myopathy

We propose that, under basal conditions, inter-domain interactions within Sis1 are maintained through transient contacts primarily mediated by the intrinsically disordered GF and GM domains (Figure 9A, black arrows). Specifically, the GM domain interacts dynamically with the C-terminal (CTD) and J-domains. In addition, a stabilizing J-CTD contact functionally masks the conserved HPD motif in the J-domain, thereby preventing unregulated recruitment of Hsp70 (Figure 9A).

**Figure 9.**
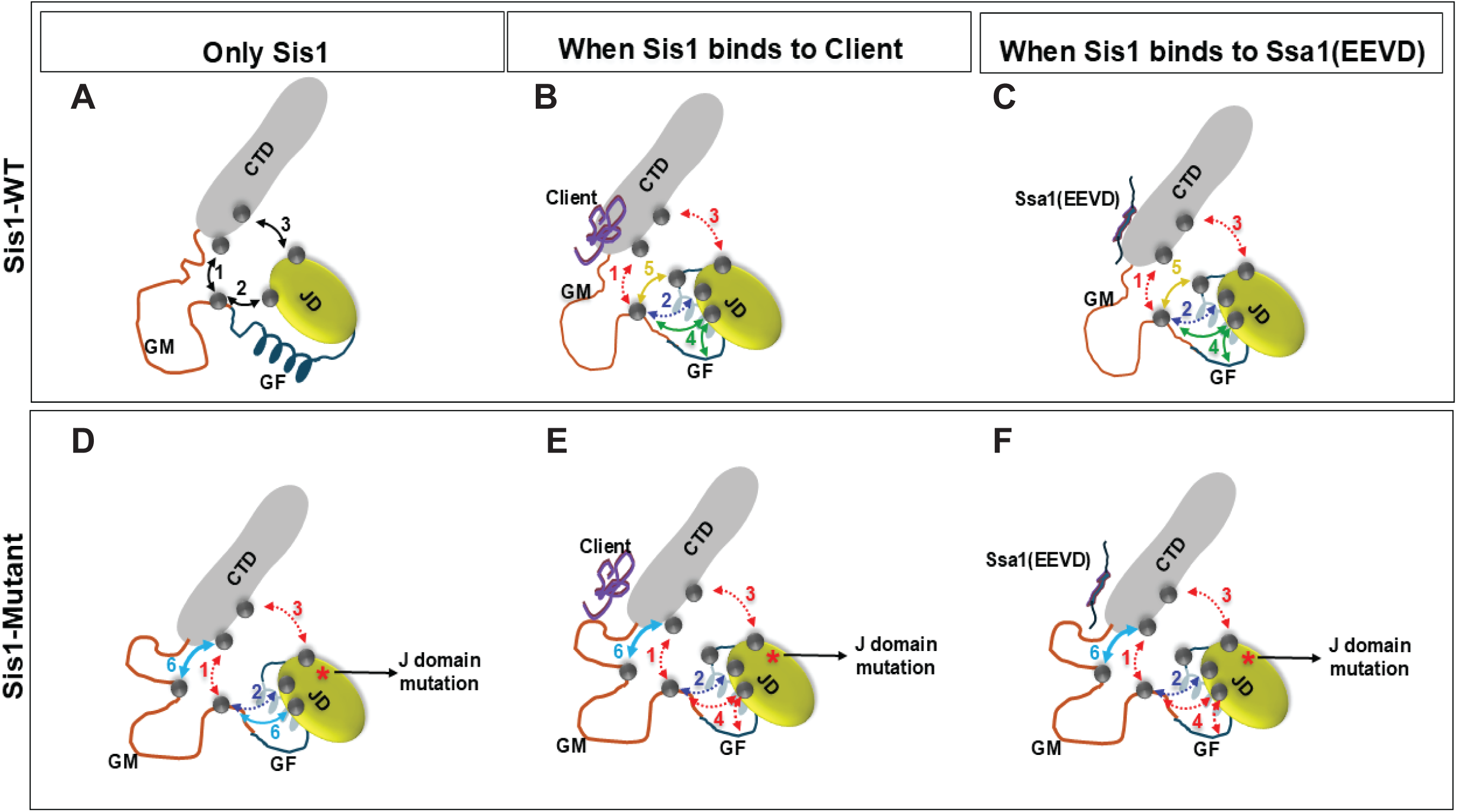
Diagrammatic representation of the proposed contacts and mechanistic features of the Hsp40-Hsp70 complex. The cartoon shows the different domains of Sis1: JD (olive), GF (dark blue), GM (orange), and CTD (grey). Grey dots represent residues that form interactions between domains. Arrows represent changes in interactions as follows: black represents interactions in Sis1-WT, red and blue represents lost and weakened interactions; respectively, green and yellow represents new acquired interactions when client and/or Ssa1-EEVD binds to Sis1, cyan represents interactions in Sis1-mutants. Numbers in black, red, and blue represent transient inherent, lost, and weakened interactions; respectively in both Sis1-WT and mutant. Numbers in green and yellow represent gained interactions formed when Sis1-WT binds to either client and/or Ssa1-EEVD. Numbers in cyan color represent interactions in mutants. When Sis1-WT binds to client and or Ssa1-EEVD transient inherent interactions of Sis1 (A) either get lost or weakened, while J-domain form new interactions with GF and GM domain (B) and (C). All the transient inherent interactions present in Sis1-WT are either lost or weakened in Sis1 J-domain mutants (D). This is accompanied by additional GM-CTD and J domain-GM interactions (D). When these mutants bind to either client and or Ssa1-EEVD, there are no new gained interactions as observed with Sis1-WT binding to either of them (E) and (F). However, the J domain-GM interaction which was there in mutant is lost with both client and/or Ssa1-EEVD engagement.

Upon engagement of the CTD with either client or the EEVD motif of Ssa1, this inhibitory J-CTD interaction, along with specific GM-CTD contacts, is lost (Figure 9B–C, red dashed arrows). Consequently, flexible regions of the GM domain reorient to establish new interactions with the GF and J-domains (Figure 9B-C, green and yellow arrows), weakening the GM-J-domain contacts observed in the unbound state (Figure 9B-C, blue arrows). This remodeling alters the arrangement of the GF-J and GF-GM interfaces (Figure 9B-C, black arrows), which expose the GF region to bind the substrate binding domain (SBD) of Ssa1, accompanied by release of autoinhibition and subsequent ATPase activity for refolding.

Interestingly, disease mutations within the J-domain impair both the J-CTD and GM-CTD interactions, suggesting allosteric communication between these regions (Figure 9D, red arrows). These mutations favor persistent GM-CTD interactions even in the presence of client or the EEVD motif (Figure 9D–F, cyan arrows). This is consistent with molecular dynamics simulations showing reduced availability of CTD residues for client or EEVD binding. In mutant complexes with either the EEVD motif or the Rnq1 peptide, key CTD residues engage non-specifically with the GM domain, occluding their functional interaction sites. These observations are further supported by inter-domain contact analysis (Supplementary Figure 9). However, the mutation-driven reduction in specific GM residues available for these transitions appears to allosterically disrupt the GF-J and GF-GM contacts (Figure 9E–F, red arrows). These altered inter-domain interactions potentially correlate with impaired binding with the client (Figures 2 and 6G) and likely underlie the observed defect in substrate refolding (Figure 1C).

Our findings suggest that J-domain mutations perturb Sis1 function through both direct and indirect mechanisms: directly, by disrupting J-GF contacts; and indirectly, by destabilizing the J-CTD inhibitory linkage. Both routes converge on similar inter-domain rearrangements, indicating a unified allosteric pathway triggered by J-domain mutation or client/Ssa1 engagement.

## DISCUSSION

One of the significant challenges in designing therapeutics for LGMDD1 is the lack of a solid mechanistic understanding of the disease’s pathogenesis. In recent years, our lab and others have proposed multiple theories regarding how mutations in the Hsp40 co-chaperone DNAJB6 affect its interaction with Hsp70 and client processing, ultimately leading to LGMDD1(*19*, *33*, *58–60*). While much of this research has focused on co-chaperone interactions, a critical question remains: How convincing are these theories in the absence of a skeletal muscle protein client? This question is particularly relevant since these mutants exhibit client-specific and conformer-specific defects. Most recent studies aiming to understand the function of DNAJB6 disease mutants have concentrated on the differences in their interaction with Hsp70 (*19*, *33*, *58*, *59*). They highlight the significance of releasing autoinhibition from the GF domain of DNAJB6 and its impact on co-chaperone function. However, it is essential to note that these studies do not involve a muscle client protein. In our previous work, we successfully demonstrated how GF mutations in a DNAJB6 homolog, Sis1 in yeast, affect chaperone activity in the presence of both client proteins and their conformers and Hsp70 (*18*, *19*). In this study, we applied a similar approach to investigate how mutations in the J-domain of DNAJB6, using its homolog Sis1, influence its functioning concerning client proteins and co-chaperones. Additionally, through an in-silico approach, we showed how these mutants differ structurally in their molecular interactions with clients and co-chaperones, further validating our functional findings. We found that the LGMDD1 J-domain mutant’s ability to stimulate Hsp70 ATPase activity was reduced as compared to the wild-type protein (Fig. 3C-D), which may result in general impairment of protein quality control and accumulation of protein inclusions in muscle. All mutants showed decreased binding to substrates (Fig.2A, B), with E53K< S49V= E53A< N56L for FL Rnq1, and E53A< S49V<E53K< N56L for luciferase as substrates. The LGMDD1 mutants showed similar binding defects to Hsp70 (N56L is worse without client) (Fig. 2E). The mutants showed a similar reduction in Hsp70 ATPase stimulation, but this was client conformation-dependent, as expected based on previous work (Fig.3A-D). Finally, the mutants showed a similar reduction in luciferase refolding efficiency (Fig. 1A). From our molecular dynamics simulation studies, we found that interactions between the J-domain and the GM and CTD domains serve as regulatory interfaces that coordinate client binding and Hsp70 activation. Mutations that disrupt these contacts impair substrate processing and co-chaperone function, highlighting their critical role in maintaining chaperone activity (Figures 4-9).

The binding of Hsp40 to its client is crucial for downstream substrate processing. Rnq1 is a well-known client of Sis1 and is extensively studied as a yeast prion protein (*48*). It has a large C-terminal region of about 250 residues that is, likely prionogenic. Mapping the actual and potential prion structures of Rnq1 in two different [*RNQ+*] variants indicates that these structures extend beyond the C-terminal core of 40 residues (*48*). Transferring a prion from the full-length Rnq1 to its truncated versions also leads to significant alterations in the prion structures (*48*). The binding interactions between Sis1 mutants and denatured full-length Rnq1, as well as its truncated form ΔN Rnq1 demonstrate varying degrees of client association as compared to Sis1-WT (Figure 2A and 6G). Jelen et al. demonstrated that precise binding of the client to Hsp40 enables effective binding of Hsp70 to the complex (*61*). When Ssa1 was introduced to either Sis1-WT or mutants complexes with denatured FLRnq1, the ATPase activity of Ssa1 remained unchanged in all instances (Figure 3A). The HPD motif of the J-domain plays a critical role in activating the ATPase activity of Ssa1. It is a pivotal switch that triggers ATPase activity, facilitating client transfer to Ssa1 and subsequent refolding. Previous studies have indicated that the HPD motif on the J-domain interacts with the binding groove of the nucleotide-binding domain (NBD) of Ssa1 (*62*, *63*). This suggests that the binding of Ssa1 to the HPD motif is likely unaffected or that mutations in the J-domain do not alter HPD’s binding to Ssa1. This was further supported by our molecular dynamics simulations of Sis1 and its mutants in complex with Ssa1-EEVD [Supplementary Figure 10]. However, the overall binding of Ssa1 to the Sis1-FLRnq1 complex varies among mutants and is reduced as compared to the wild-type (Figure 2D).

Ssa1 binds to multiple sites on Sis1. In addition to the HPD motif of the J-domain, which activates Ssa1’s ATPase activity, the EEVD motif located at the C-terminus of Ssa1 is known to be important for binding to Sis1 and facilitating client refolding (*57*). As shown previously, the EEVD motif binds to lysine residues on the C-terminal domain (*53*, *57*). Recent studies have indicated that Ssa1 competes with clients for binding to these client-binding residues. Our molecular dynamics simulation studies suggest that the Ssa1-EEVD binding to Sis1 is reduced in mutants. The experimentally observed reduction in Ssa1 binding may also be attributable to decreased binding at other sites on Ssa1. In Sis1 homologs, the EEVD motif induces a conformational change in the GF domain, which releases J-GF domain auto-inhibition, thereby allowing the sequestered HPD motif on the J-domain to be available for additional Ssa1 binding (*27*). Reduced binding of Ssa1-EEVD could drive changes in distant J-GF interactions, affecting the binding of SBD and NBD of Ssa1 on Sis1, ATPase activity and subsequent refolding.

It has been reported that DnaJ remodels misfolded proteins through a conformational selection mechanism, while DnaK induces conformational expansion of the client protein via mechanical unfolding (*64*). This study also provided direct evidence that the protein client undergoes repeated cycles of binding and release from chaperones during refolding, and it was observed that the rate of chaperone cycling correlates with an improved refolding yield (*64*).

Recently, research on auto-inhibited DNAJB6 constructs has shown that, in combination with the J-domain, the GF-linker (both G-rich region and Fx repeats of H5 helix) creates a hydrophobic, partially collapsed cluster that exhibits a notable degree of allosteric communication. Disruption of this communication can destabilize autoinhibition (*47*), indicating the importance of the G-rich region in driving autoinhibition. Beyond its intramolecular role, the GF-linker can be recognized by the substrate-binding domain of Hsc70, influencing the lifetime of the entire JDP-Hsc70 complex. Notably, both intra- and intermolecular functions of the GF-linker are specific to the DNAJB class members (*47*). Our results show that distant EEVD and client engagement with Sis1 drive the interaction of J and GM domain with the G-rich region of the GF linker (Supplementary Figures 11 and 12). In the J-domain mutants, distant interactions are disrupted, and unregulated new interactions occur, and as such, the G-rich region binding to the J and GM domains is strikingly reduced (Figure 8). This suggests that overall decreased binding of Sis1 mutants to client or Ssa1 could also be due to loss of the G-rich region contacts.

In humans, Hsp40 proteins can function as monomers, dimers, or oligomers (*46*). Classical Hsp40 members assemble into homodimers or mixed J-protein complexes through conserved C-terminal motifs, binding unfolded substrates through their conserved β-barrel C-terminal domains (CTDs) (*26*, *65*, *66*). A subset of nonclassical Hsp40s, such as DnaJB2, DnaJB6b, DnaJB7, and DnaJB8, have a CTD domain architecture that differs from the classical dimeric DnaJ orthologs (*14*, *26*, *58*, *66*, *67*). Notably, DNAJB8 and DNAJB6b self-assemble into soluble oligomers both in vitro and in vivo (*14*, *68*, *69*). A recent study of DNAJB8 found that the interactions between the J-domain and CTD regulate the recruitment of Hsp70, indicating a built-in regulatory mechanism that controls the recruitment and activation of Hsp70 (*46*). We found that there is a crucial interaction between the J-domain and the CTD domain of Sis1 (Supplementary Figures 5 and 9) that gets lost with either the client or the Ssa1-EEVD binding (Figures 5 and 9), suggesting a similar regulatory mechanism.

The degree of homology between Sis1 and DNAJB6 and how our findings with Sis1 relate to the disease mechanism is a valid question. While we acknowledge some concerns and recognize there are conflicting reports regarding Sis1 as a homolog of DNAJB6, it is essential to note that Sis1 is more homologous to human DNAJB1 than DNAJB6. However, even DNAJB1, which is assumed to be the most appropriate homolog of Sis1, differs in the GF domain region, making it challenging to estimate the degree of J-domain release in Sis1 variants based on DNAJB1. Structurally, although the number of domains differs between DNAJB6 and Sis1, there is a significant resemblance in the orientation of their J-domain and GF domain (*47*). Importantly, LGMDD1 disease mutants have been reported exclusively in these two domains. Therefore, studying Sis1 in our model system is highly relevant, especially given the absence of a client protein associated with DNAJB6 in muscle pathology. Moreover, recent studies seeking to understand the function of DNAJB6 using other substrates, such as the Huntingtin protein (polyQ), also utilize Sis1 in yeast systems, albeit with caution. We have successfully translated our findings from yeast Sis1 (Hsp40) to gain insights into muscle chaperonopathy related to the DNAJB6 protein through our previous studies (*18*, *33*). In the absence of a DNAJB6 muscle client protein, it is imperative that we utilize all available tools and systems to dissect the LGMDD1 disease mechanism. Thus far, our approach to using Sis1 as a homolog of DNAJB6 and its known clients has provided important clues needed to develop therapeutic strategies for LGMDD1. We showed previously that the dynamics of the DNAJB6-Hsp70 complex are dysregulated in muscle expressing a LGMDD1 mutant and that restoring the dynamics of the complex correlated with improved muscle strength and myopathology (JCI REF). Therefore, understanding all the structural changes induced by LGMDD1 mutants is key to identifying precisely what could be targeted for therapeutics. From this work, we believe that targeting the inter-domain network or stabilizing the GF region interaction could potentially be an important therapeutic avenue to explore for LGMDD1 myopathy.

## MATERIALS AND METHODS

### Cloning, expression, and purification of recombinant proteins

Sis1-WT, Sis1 mutants (S49V, E53A, E53K, N56L), and Ssa1-WT were cloned into the pPROEx-Htb vector obtained from Addgene. This plasmid encodes a hexa-His-tag, a TEV cleavage site, and the respective cloned gene for expression. The Sis1 mutants were generated using the Quick-Change Mutagenesis Kit from Agilent Technologies (200517), with primer sequences designed using Agilent’s online primer design program. The mutations were confirmed by sequencing the entire coding region of Sis1. Sis1-WT and the mutants were expressed at 16 °C, while Ssa1-WT was expressed at 18 °C to enhance the yield of soluble protein. All purification steps were conducted at 4 °C. Protein purity exceeded 99%, as determined by SDS/PAGE and Coomassie staining. The final protein concentration was estimated using the Bradford assay, with bovine serum albumin as the standard. After purification, all proteins were frozen in liquid nitrogen and stored at −80 °C until further use.

The purification of Sis1-WT and the mutants followed a standard protocol with some modifications. The proteins were isolated from *Escherichia coli* strain Lemo 21(DE3) (New England Biolabs #C2528H) grown in 2X YT medium at 30 °C until the optical density (OD600) reached 0.6–0.8. Cultures were induced with 0.5 mM IPTG and incubated overnight at 16 °C. Cells were harvested and lysed in buffer A (50 mM sodium phosphate buffer, pH 7.4; 300 mM NaCl; 5 mM MgCl2; 10 mM imidazole; 0.1% Igepal; 0.01 M TCEP (tris(2-carboxyethyl)phosphine) (Thermo-Fisher Scientific #20491), along with a protease inhibitor cocktail (EDTA-free) (Thermo-Fisher Scientific #A32965) and a pinch of DNase I (Millipore Sigma #DN25). Following cell lysis, debris was removed via centrifugation at 20,000 × g, and the supernatant was applied to cobalt-based Talon metal affinity resin (Thermo-Fisher Scientific #89964). After washing, proteins were eluted in gradient fractions using buffer A containing increasing imidazole concentrations (150–400 mM) (Millipore Sigma #I0125). Purified proteins were treated with His-TEV protease (purified in the lab) at 30 °C for 1 hour, extensively dialyzed at 4 °C, and passed through Talon metal affinity resin again to remove the cleaved His tag and His-TEV protease. The resulting pure proteins were concentrated and stored at −80 °C.

For Ssa1-WT purification, a similar established protocol was used. Proteins were isolated from *Escherichia coli* strain Rosetta 2(DE3) (Novagen, EMD-Millipore #71397-3) and cultured in LB medium with 300 mM NaCl at 30 °C until the OD600 reached 0.6–0.8. Protein expression was induced with 0.5 mM IPTG added to the culture and incubated overnight at 18 °C. Cells underwent lysis in buffer A (20 mM HEPES, 150 mM NaCl, 20 mM MgCl2, 20 mM KCl), supplemented with a protease inhibitor cocktail (EDTA-free) using lysozyme (Millipore Sigma #L6873). The remaining purification steps were analogous to those for Sis1-WT and its mutants, except Ssa1-WT were eluted with buffer A containing 250 mM imidazole. The full-length Rnq1-WT, and ΔN Rnq1 proteins were purified in denatured form following a previously published method from our laboratory (*70*).

### Kinetic Assay

Kinetic assays for fiber formation were conducted using a SpectraMax M3 fluorimeter microplate reader. The Rnq1 monomer was diluted using a 7 M guanidine hydrochloride to a final concentration of 4 μM in Fiber Formation Buffer (FFB), which contains 150 mM NaCl, 5 mM KPO4, and 2 M urea at pH 6. To initiate fiber formation, a 50-fold molar excess of Thioflavin-T was added in the presence of glass beads to facilitate agitation (*70*). The change in Thioflavin-T fluorescence over time was measured with an excitation wavelength of 450 nm and an emission wavelength of 481 nm. The plate was agitated every minute before reading for 10 seconds.

#### Luciferase refolding assay

Heat-denatured refolding of luciferase was conducted as previously detailed (*71*). In brief, Ssa1 (2 µM) was incubated in a refolding buffer of 50 mM Tris (pH 7.4), 150 mM KCl, and 5 mM MgCl2. This buffer was supplemented with 1 mM ATP and an ATP regenerating system, which included

10 mM phosphocreatine and 100 mg/mL phosphocreatine kinase, for 15 minutes at room temperature. Following this incubation, luciferase (25 nM) was added, and the mixture was further incubated for 10 minutes. Subsequently, Sis1-WT/mutants (0.05 µM) were introduced, and the mixture was subjected to a heat shock at 44 °C for 20 minutes. The reactions were then rapidly returned to room temperature. Finally, 25 µL aliquots of the refolding mixture were obtained and combined with 50 µL of luciferase assay reagent from Promega Corporation. At designated time points, the enzymatic activity was measured using a SpectraMax M3 luminometer microplate reader (Molecular Devices).

### Binding Assays

#### Substrate-binding ELISA assays

The substrate-binding ELISA assays were conducted with modifications to previously established methodologies (*31*). Three distinct substrate proteins— Rnq1, ΔN Rnq1, and firefly luciferase (Promega Corporation)- were denaturized for one hour at a temperature of 25 °C in a buffer consisting of 3 M guanidine hydrochloride, 25 mM HEPES (pH 7.5), 50 mM KCl, 5 mM MgCl2, and 5 mM DTT. Following the denaturation process, the substrates were diluted in 0.1 M NaHCO3 and subsequently bound to the wells of microtiter plates (CoStar 3590 EIA plates, Corning) at concentrations of 0.4 µg for Rnq1 and 0.1 µg for luciferase. The excess unbound substrate was removed thoroughly with phosphate-buffered saline (PBS). To block unreacted sites, 100 µl of 0.2 M glycine was added for 30 minutes at room temperature, followed by additional washing with PBS-T, which contains 0.05% Tween 20. The wells were treated with 0.5% fatty-acid-free bovine serum albumin (BSA) (Millipore Sigma) in PBS to further diminish nonspecific binding for six hours. After this blocking step, the wells were washed once more with PBS-T. Sis1-WT and various Sis1 mutants were serially diluted in PBS-T supplemented with 0.5% BSA and incubated with the substrates overnight at room temperature. Following extensive washing with PBS-T to eliminate any unbound proteins, rabbit anti-Sis1 antibody (CosmoBio) was introduced at a dilution of (1:15000) and incubated for two hours at room temperature. The process continued with further washings before the addition of donkey anti-rabbit HRP-conjugated secondary antibody (Millipore Sigma, diluted 1:4000). The amount of Sis1 retained in each well was quantified using a tetramethyl benzidine/H2O2 (TMB peroxidase EIA substrate) kit (Bio-Rad). The resultant color intensity was then measured at a wavelength of 450 nm using a SpectraMax M3 microplate reader, with the reaction terminated by the addition of 0.02 N H2SO4.

#### Ssa1-binding assays

In the Ssa1-binding assays, Ssa1 was immobilized at a concentration of 200 nM onto the wells of a microtiter plate. Subsequently, various dilutions of purified Sis1-WT and Sis1 mutants were introduced for incubation. The quantity of bound Sis1 was detected following the previously outlined methodology, employing a SpectraMax M3 microplate reader for measurement.

#### Substrate bound Sis1 combined with Ssa1 binding assays

Denatured substrates (Rnq1 and luciferase) were mixed with Sis1-WT and Sis1 mutant proteins at a concentration of 5 nM. This mixture was incubated at room temperature for 1 h and adsorbed onto the microtiter-well plates. Following incubation and extensive washing, serial dilutions of Ssa1-WT were introduced into the wells. The detection of bound Sis1 was achieved through probing with a mouse-Hsp70 (Ssa1) antibody at a dilution of 1:2000 (Abcam), followed by the application of a goat anti-mouse IgG [H+L] HRP-conjugated secondary antibody (Thermo-Fisher Scientific). Quantification of Sis1 was carried out using a SpectraMax M3 microplate reader.

### Colorimetric determination of ATPase activity

The ATPase activity was assessed utilizing a refined assay protocol based on previously established methods (*72*). The ATPase reagent was prepared daily, comprising 0.081% W/V Malachite Green, 2.3% W/V polyvinyl alcohol, and ammonium heptamolybdate in 6 M HCl in a specific ratio, allowing a stable green/golden solution to form after standing for 2 hours. Before utilization, this reagent was filtered through 0.45 μm syringe filters (Millipore Sigma). ATPase activity was initially evaluated in the absence of any client protein by incubating Sis1-WT/mutants and Ssa1-WT in varying concentrations (0.05:1.0 µM) with 1 mM ATP in a defined assay buffer, which consisted of 0.02% Triton X-100, 40 mM Tris–HCl, 175 mM NaCl, and 5 mM MgCl2 (pH 7.5) at 37 °C for designated time intervals, as indicated in the figure legends. Upon incubation, 25 µL of the reaction mixture was collected and placed into wells of a 96-well plate, followed by the addition of 800 µL of the ATPase reagent and 100 µL of 34% sodium citrate to halt ATP hydrolysis. After a 30-minute incubation at room temperature, absorbance was measured at 620 nm using a SpectraMax M3 fluorimeter microplate reader. A control sample containing ATP alone in the buffer was similarly processed to correct any intrinsic ATP hydrolysis. Furthermore, a phosphate standard curve was established daily using potassium phosphate to ensure measurement accuracy. For ATPase activity assays involving client proteins, the same methodological approach was employed, incorporating client protein Rnq1 monomer (25 µM) and Rnq1 amyloid (10% of the final reaction mixture), with the amyloid formed at temperatures of 18°C, 25°C, and 37°C to examine their effects on chaperone activity and ATP utilization.

### System Preparation

All molecular dynamics (MD) simulations were performed using GROMACS (version 2020.4). The system was prepared by solvating the protein in a periodic cubic box with explicit solvent molecules, followed by the addition of counterions to maintain charge neutrality. The force field used for all-atom simulations was CHARMM36m (*45*, *73*).

### All-Atom Molecular Dynamics Simulations

For all-atom simulations, energy minimization was carried out using the steepest descent algorithm to remove steric clashes and ensure system stability before equilibration. The minimization was performed for 50,000 steps with an energy convergence tolerance (emtol) of 1000.0 kJ mol−1 nm−1. Position restraints were applied to both the backbone and side-chain atoms using harmonic force constants of 400 kJ mol−1 nm−2 and 40 kJ mol−1 nm−2, respectively. A Verlet neighbor list was used with an update frequency of 10 steps. The van der Waals (vdW) interactions were modeled using a cut-off scheme with a force-switch modifier, where interactions were smoothly switched off between 1.0 nm and 1.2 nm. Electrostatic interactions were treated using Particle Mesh Ewald (PME) with a real-space cut-off of 1.2 nm. Hydrogen bonds were constrained using the LINCS algorithm. Following minimization, the system was equilibrated using two phases of NVT and NPT simulations at 310 K. Equilibration was conducted using a time step of 1 fs for a total of 1.25 ns (1,250,000 steps). Temperature coupling was applied using the velocity-rescale (v-rescale) thermostat with a coupling time constant (τT) of 1.0 ps, maintaining separate temperature groups for the solute (SOLU) and solvent (SOLV). The neighbor list update frequency was set to every 20 steps, with a vander Waals cut-off of 1.2 nm and PME electrostatics as in the minimization stage. Position restraints were maintained on the solute backbone and side chains during equilibration. For efficient center-of-mass motion removal, linear momentum removal was performed every 100 steps for both SOLU and SOLV. Initial velocities were generated from a Maxwell-Boltzmann distribution at 310 K. MD runs were performed in the NPT ensemble for 10 ns (2,500,000 steps) with a time step of 4 fs. The temperature was maintained at 310 K using the v-rescale thermostat with separate coupling groups for solute and solvent. The Parrinello-Rahman barostat was used for pressure coupling, maintaining an isotropic pressure of 1.0 bar with a compressibility of 4.5 × 10−5 bar−1 and a pressure coupling time constant (τP) of 5 ps. All energy minimization, equilibration and MD runs were performed using the GROMACS module on the Bridges2 server (*74*).

### Molecular Dynamics Trajectory Preparation and Parsing

Atomistic molecular dynamics (MD) simulations were conducted for full-length wild-type and mutant variants of the protein under multiple experimental scenarios, including ligand-bound and unbound forms, peptide interactions, and domain truncations. Trajectories were analyzed using Python (v3.10) with the MDAnalysis, MDTraj, and NumPy packages. Simulations were preprocessed by stripping solvent and ions, and only protein atoms were retained for downstream structural analysis. In all cases, simulation time was discretized into 10,000–50,000 frames, depending on system size and trajectory length, ensuring temporal resolution sufficient for contact-based statistical analysis. Global and local structural compactness and flexibility were evaluated using radius of gyration (Rg) metrics. Rg was computed as the mass-weighted mean square distance from the center of mass for all protein atoms across the trajectory. The outputs were smoothed using rolling averages over 50-frame windows and plotted to identify regions of differential flexibility or compaction across conditions.

### Residue–Residue Contact Analysis

Cα–Cα or heavy-atom distance matrices were computed for all residue pairs across each frame using a sliding distance threshold (typically ≤ 8.0 Å for Cα–Cα and ≤ 4.5 Å for heavy-atom contacts) by using Conan 1.1 software package (*75–77*). A contact probability matrix was generated for each system by averaging over all frames. In analyses comparing multiple conditions (e.g., WT vs. mutant), the contact probability matrices were subtracted to yield a differential contact map. Only differences exceeding a predefined threshold (|ΔP| ≥ 0.2) were considered significant and retained for visualization. Domain-wise contact matrices were computed by grouping residues into functional domains using a user-defined mapping dictionary, summing all pairwise contacts between domains, and normalizing by total frame count.

### Gain/Loss Contact Identification and Visualization

To quantify residue pairs exhibiting condition-specific contact formation, a gained/lost contact map (difference contact map) was constructed by filtering the differential contact matrix based on two-sided thresholds (gain: ΔP > +0.3, loss: ΔP < -0.3). These were visualized using diverging colormaps (red for loss, blue for gain) in matplotlib to emphasize structural reorganization between conditions. For ease of interpretation, contact matrices were optionally symmetrized, and domain annotations were overlaid along the matrix axes.

### Kullback-Leibler Divergence (KLD) for Structural Ensemble Comparison

To quantify ensemble differences between mutant variants, the Kullback–Leibler Divergence (KLD) was calculated on per-residue contact probability distributions. Contact probability vectors for each residue were normalized across all conditions. The KLD metric was then used to detect residues whose local contact environments diverged significantly between variants, highlighting functionally important or conformationally distinct sites. KLD profiles were plotted along the protein sequence, and peaks exceeding 0.1 bits were interpreted as structurally differentiating regions.

### Per-Residue Contact Frequency Profiles

Each residue’s interaction propensity was quantified by calculating the mean number of contacts formed per frame, averaged across the trajectory. Contact definitions used a Cα–Cα cutoff of ≤ 8 Å, and frequencies were normalized to a per-frame basis. These contact frequency profiles were compared across conditions (e.g., mutant vs. WT) to identify residues exhibiting significantly altered local interaction densities. Differences were plotted as line plots or bar graphs, and statistical significance (e.g., t-tests across frames) was optionally applied.

### Peptide and EEVD Binding Analysis

Contact interfaces between the protein and peptide ligands (including the EEVD motif) were characterized using residue-wise interaction frequency calculations. Any contact between the peptide and the protein was counted if any heavy atom from each residue pair came within 4.5 Å during a given frame. Contact frequencies were computed for each residue pair, and a peptide contact profile was generated by summing across all peptide residues for each protein residue. Comparative peptide-binding maps were then generated for wild-type and mutant forms, and the differences were visualized as shaded plots highlighting shifted interaction hotspots.

### Statistical and Computational Implementation

All analyses were performed using Python 3.10 with custom scripts leveraging the MDAnalysis (v2.3.0), MDTraj (v1.9.7), matplotlib (v3.7), and seaborn (v0.12) libraries on Bridges2 server (*74*). Data aggregation and statistical operations (e.g., moving averages, per-frame normalization, filtering by threshold) were performed using NumPy and pandas. Heatmaps were generated with fixed color scales to enable cross-condition comparisons. For all differential analyses, contacts or metrics were considered significant if their difference exceeded pre-defined thresholds and persisted across >50% of the trajectory frames.

## Acknowledgments

We are thankful to the True Lab and the Weihl Lab members for helpful discussions and comments on the manuscript. We would also like to thank Sun Joo Lee and Adam Balutowski for their initial help with molecular simulations.

## Author contributions

Conceptualization: AKB, HLT; Methodology: AKB, GA, AJ, DC, JD; Investigation: AKB, GA, AJ; Supervision: HLT; Writing-original draft: AKB, GA, AJ, HLT; Writing-review & editing: AKB, GA, AJ, CCW, HLT.

## Data and materials availability

All data needed to evaluate the conclusions in the paper are present in the paper and/or the Supplementary Materials. Further information and requests for resources and reagents should be directed to and will be fulfilled by the lead contact, HLT.

