## Supplementary Figures for "Mutations in Hsp40 co-chaperone change the unique canonical inter-domain interactions stimulating LGMDD1 myopathy"

### Supplementary Table 1

| | | RMSD (Ref-J-domain,4rwu) | RMSD (Ref-CTD, 13cg) |
|---|---|---|---|
| AlphaFold Published | Model | 0.309 | 0.543 |
| AlphaFold3 | Model0 | 0.268 | 0.784 |
| | Model1 | 0.294 | 0.821 |
| | Model2 | 0.274 | 0.994 |
| | Model3 | 0.26 | 0.852 |
| | Model4 | 0.27 | 0.749 |
| Robetta | Model0 | 0.522 | 1.471 |
| | Model1 | 0.564 | 1.413 |
| | Model2 | 0.625 | 1.407 |
| | Model3 | 0.625 | 1.39 |
| | Model4 | 0.694 | 1.444 |

**Supplementary Table 1.** Backbone RMSD values of homology models of Sis1WT using the crystal structure of the J-domain and the crystal structure of the Peptide binding domain as a reference.

### Supplementary Figure 1

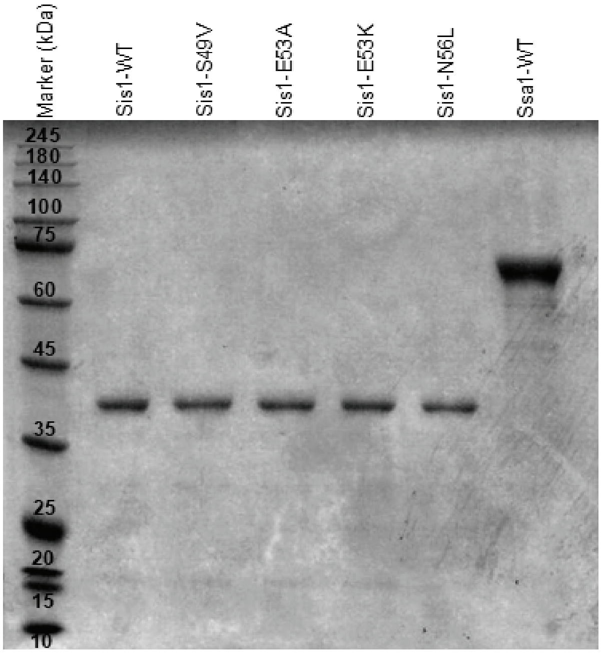


**Supplementary Figure 1. SDS-PAGE of Sis1-WT and mutants.** SDS-PAGE showing the image of purified protein bands for Marker, Sis1-WT, Sis1-S49V, Sis1-E53A, Sis1-E53K, Sis1-N56L, Ssa1-WT (from left to right) used in assays. The gel was stained using Coomassie Brilliant Blue, followed by destaining and scanning the stained gel for protein bands. One representative image from n=3 is shown here**.**

#### Supplementary Figure 2

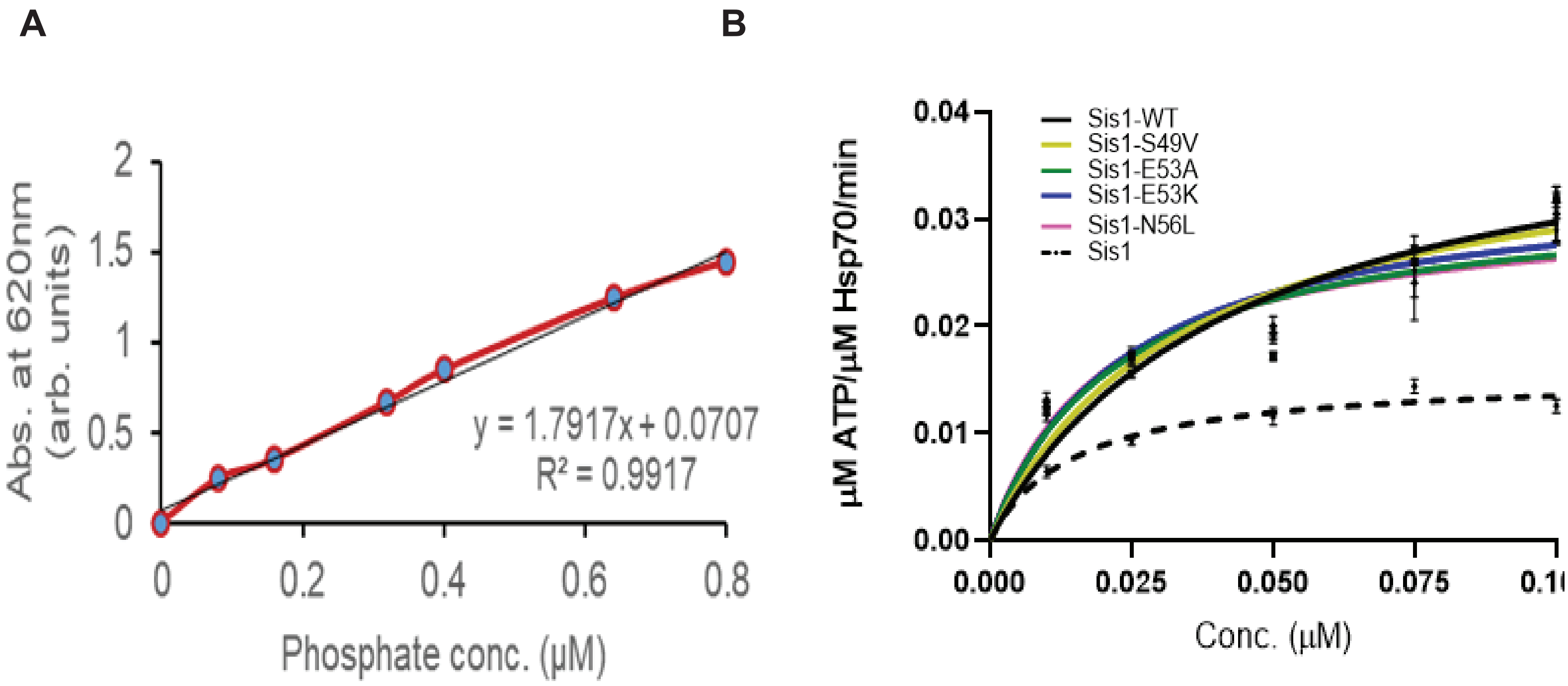


**Supplementary Figure 2.** (A) Representative Phosphate Standard Curve. ATPase assay in the absence of client protein (B). For (B), varying concentration of Sis1-WT (0.01µM-0.1µM) was incubated with Ssa1 (1µM) and ATP (1mM). Data represented as mean ± SEM, n=3 biologically independent samples. Sis1-WT was compared with LGMDD1 mutants. The differences were non-significant (NS) and are reported for the unpaired, two-sided t-test.

#### Supplementary Figure 3

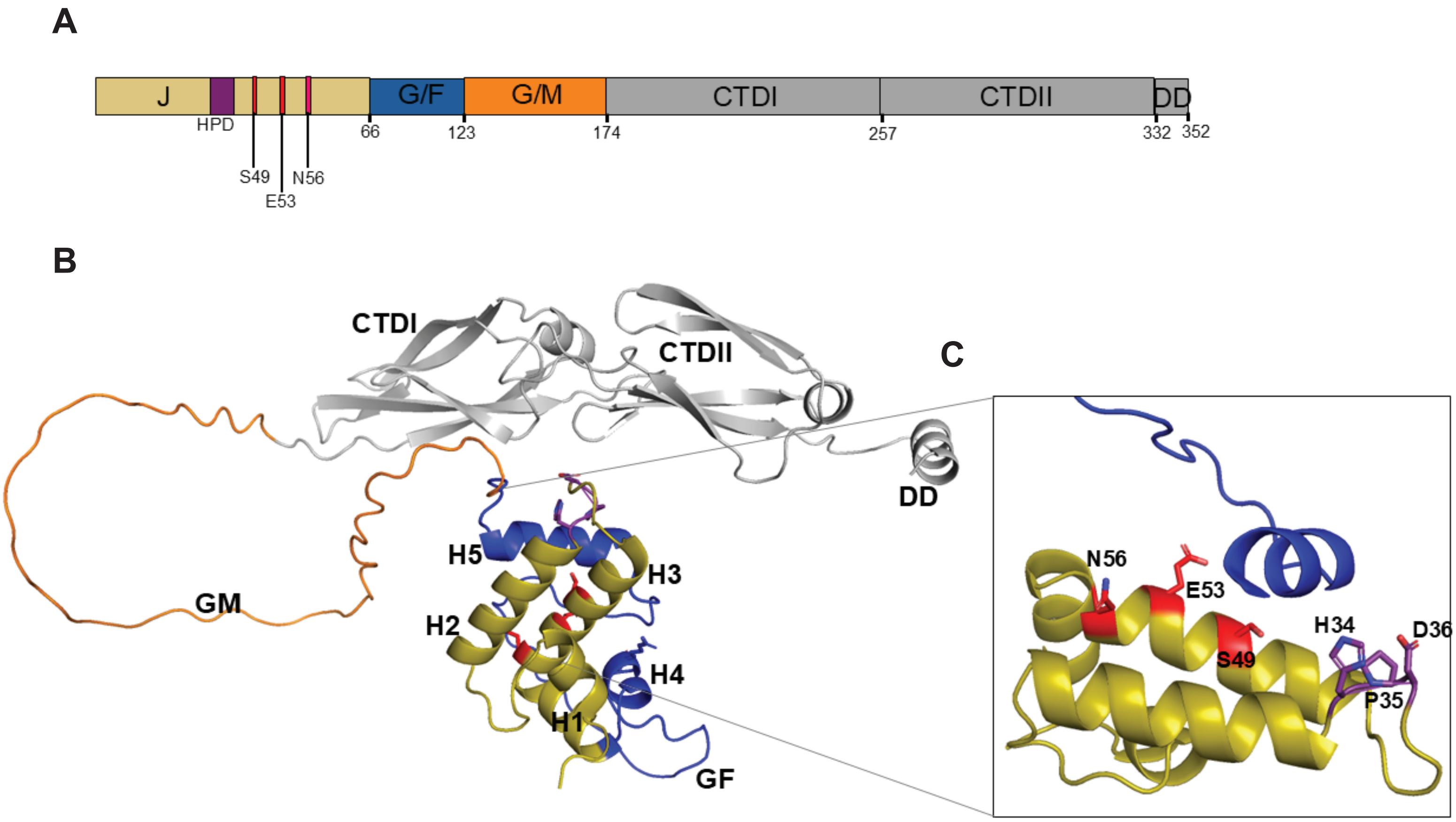


**Supplementary Figure 3. Predicted Structure of full-length Sis1WT**. (A) Schematic of the full-length Sis1WT primary sequence annotated with functional domains. Residues comprising the J-domain (residues 1–75) are indicated in olive, with point mutations S49, E53, and N56 labeled in red. The conserved HPD motif (residues 31–33) is highlighted in purple. (B) AlphaFold 3-predicted structure of Sis1WT rendered in cartoon representation. Structural domains are color-coded: J-domain (olive), GF domain (dark blue), GM domain (orange), and CTD (grey). Subdomains CTDI and CTDII are labeled; J-domain helices (H1–H4) and Helix 5 (H5) in the GF domain are indicated. (C) Enlarged view of the J-domain highlighting the positions of S49, E53, and N56 mutations (shown as red sticks) and the HPD motif residues (purple sticks) positioned on helices H2 and H3.

### Supplementary Figure 4

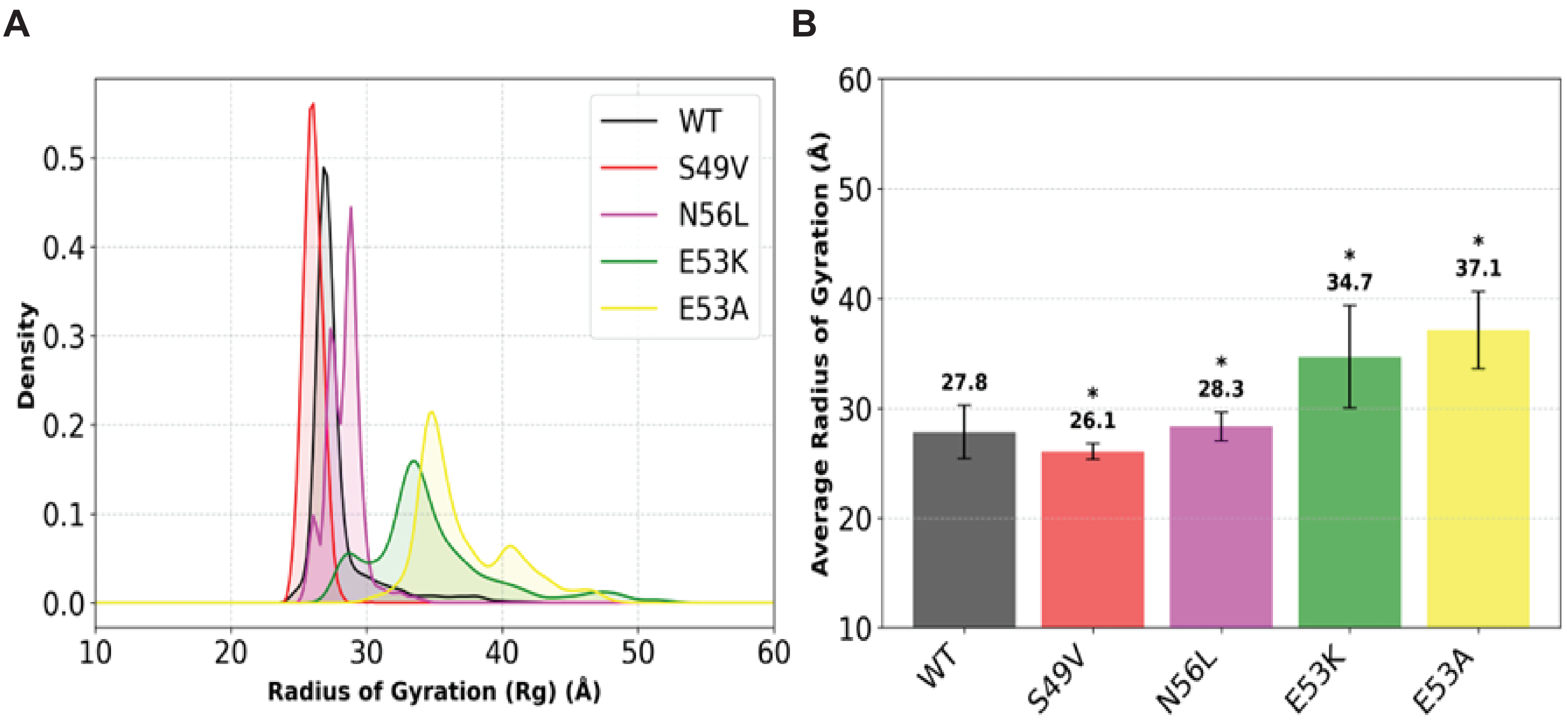


**Supplementary Figure 4. J-domain mutations modulate global compaction and conformational heterogeneity in Sis1.** Radius of gyration (Rg) profiles for backbone Cα atoms were calculated over 1000 ns MD simulations. Rg quantifies the spatial extent of atomic dispersion around the protein's center of mass. (A) Rg histograms illustrating ensemble width. Overlay of histograms of conformation ensembles of Sis1WT and mutants shows that compaction in S49V/N56L reduces conformational diversity, while E53A/E53K mutations broaden the conformational space. The intrinsic flexibility of the GF (residues 67–122) and GM (123–174) domains underlies ensemble expansion. (B) Mutants S49V and N56L (red and magenta) show reduced Rg values, indicative of a more compact ensemble relative to Sis1WT (black), while E53A and E53K mutants (yellow and green) exhibit enhanced fluctuations.

### SupplementaryFigure 5

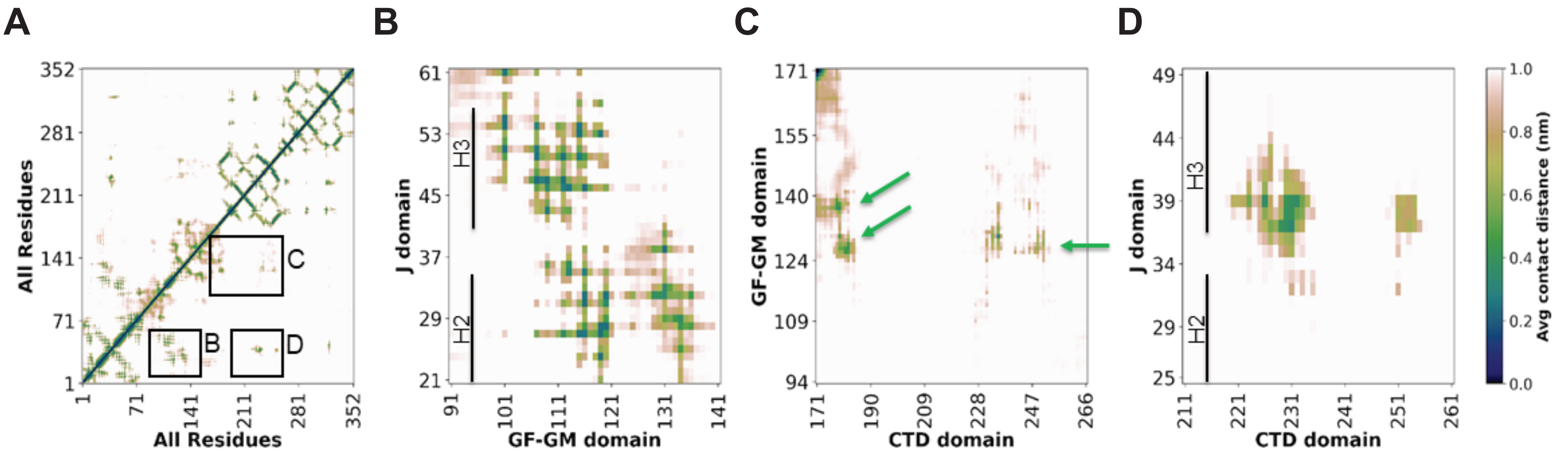


**Supplementary Figure 5. Domain-level interaction persistence in Sis1WT reveals long-range coupling across structural regions.** Average contact maps derived from a 1000 ns MD trajectory of full-length Sis1WT were generated using Conan 1.1. (A) Global contact map indicating persistent inter-residue contacts. Intensity scale reflects contact frequency; darker hues indicate more persistent interactions. Black boxes demarcate key interdomain interactions. (B-D) Zoomed-in representation of the average contacts between J-GF, GM-CTD, and J-CTD domains, respectively. (B) Close-up of the J–GF domain contacts show a robust interface where H2 helix (19-32) interacts with GF and GM domains; however, H3 helix (42-56) interacts with only the GF domain. (C) Two segments of the GM domain (one residue stretch 125-128 and the other residue stretch 130-138; shown with green arrows) interact with the CTD domain. (D) A region of the J domain in H2 helix proximal to the HPD motif forms interactions with the CTD domain. These J–CTD contacts suggest long-range coupling within the protein.

#### Supplementary Figure 6

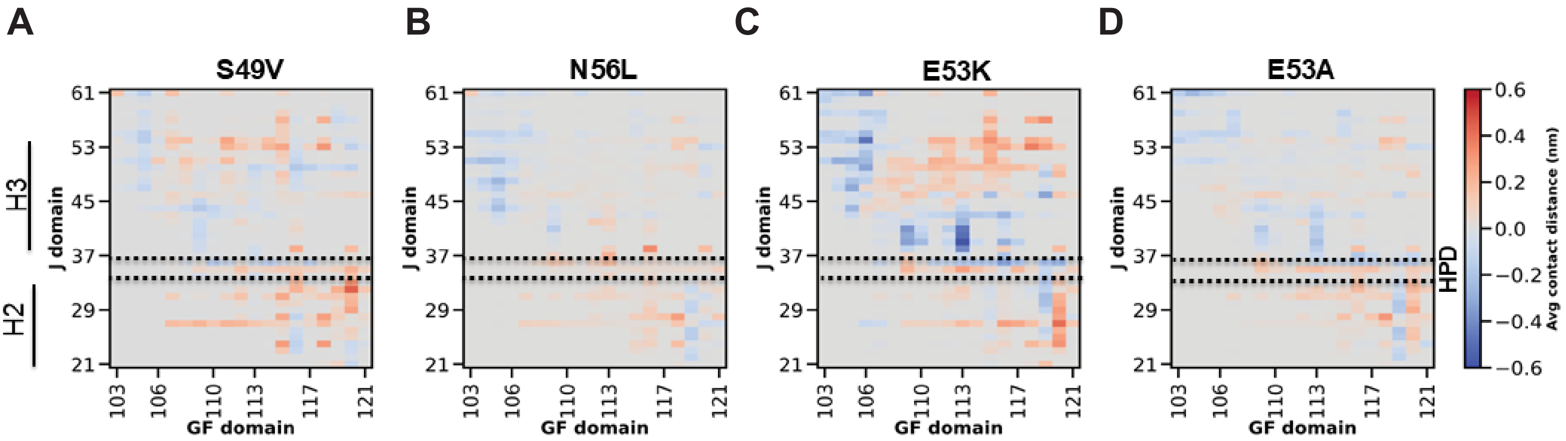


**Supplementary Figure 6. Mutations at HPD-adjacent residues uniformly reduce HPD–GF contacts without client proteins or Ssa1. T**he contacts of WT were subtracted from those of the mutants, and the difference-contact maps were plotted as the average contact distance for different domains. Positive values reflect contact loss (red); negative values indicate the formation of new interactions (blue). Difference-contact maps reveal a consistent loss of contact between the HPD motif (residues 34–36) and the GF domain across S49V (A), N56L (B), E53A (C), and E53K (D) mutants. The region of the HPD motif is marked with a horizontal dashed line. Contact maps were computed using Conan 1.1.

#### Supplementary Figure 7

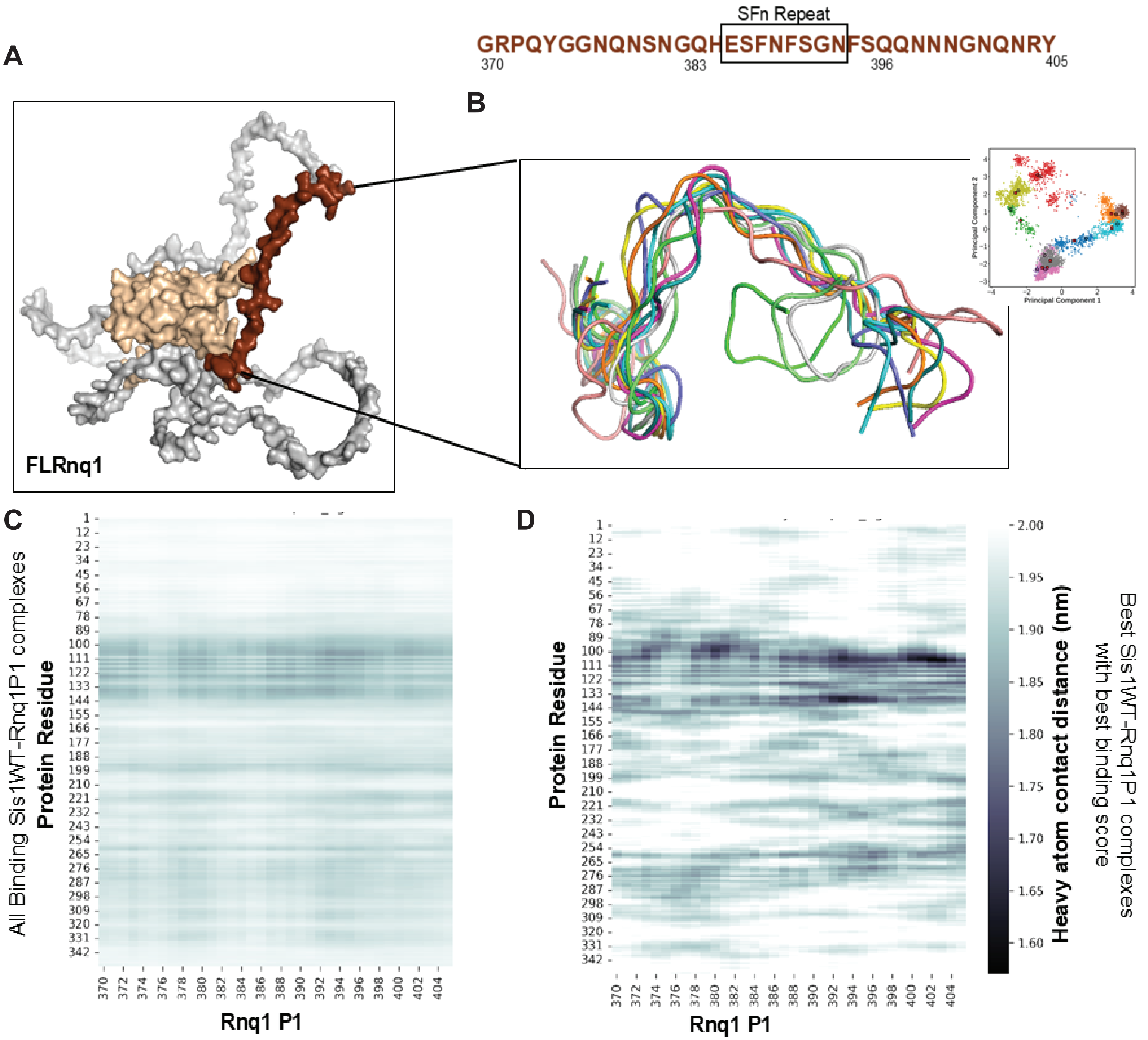


**Supplementary Figure 7. Conformational ensemble of truncated Rnq11 (Rnq1P1) reveals flexible binding modes to Sis1WT.** (A) AlphaFold structure of FLRNQ1 (AlphaFold database: AF-P25367-F1-model_v4) with Rnq1P1 highlighted in dark brown; the N-terminal domain in light brown, and the C-terminal domain in grey. (B) MD-derived conformational ensemble of Rnq1P1 clustered using dihedral Principal Component analysis. The top 10 clusters are labeled in the inset; representative conformations are circled. (C) Docking of clustered Rnq1P1 conformers to Sis1WT. Residue-resolved contact maps highlight interaction hotspots for each docked complex, revealing preferences in binding orientation.

#### Supplementary Figure 8

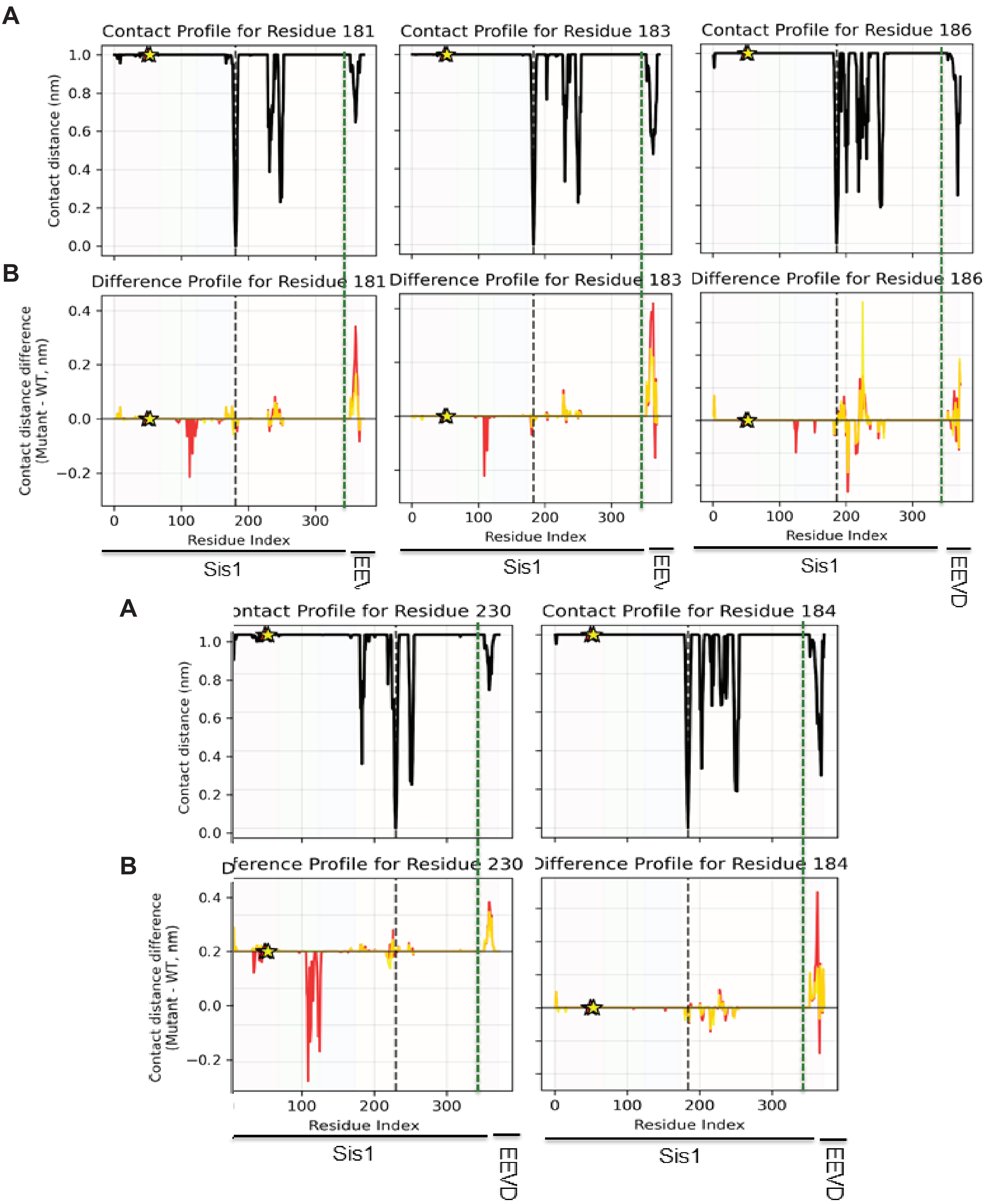


**Supplementary Figure 8. Specific CTD residues (181–253) exhibit reduced Ssa1 (EEVD) binding in Sis1 mutants.** (A) The per-residue contact profile of specific CTD residues in the Sis1WT- Ssa (EEVD) complex shows that these CTD-specific residues bind to the EEVD motif and within the CTD. The vertical black dashed line represents the position of specific CTD residues, and the vertical green dashed line represents the position from where EEVD residues start in the complex. (B) The difference in per-residue contact profiles was calculated by subtracting the average per-residue contact profile of Sis1WT from the mutants, such that positive value indicates weaker binding affinity and negative difference values indicate stronger binding than Sis1WT. Per-residue contact profiles show decreased EEVD interaction in mutants S49V (red) and E53A (yellow) for residues 181, 183, 186, 230, 250, and 253. Additionally, these residues form new contacts with other domains of Sis1. The mutated position is highlighted with * asterisk.

#### Supplementary Figure 9

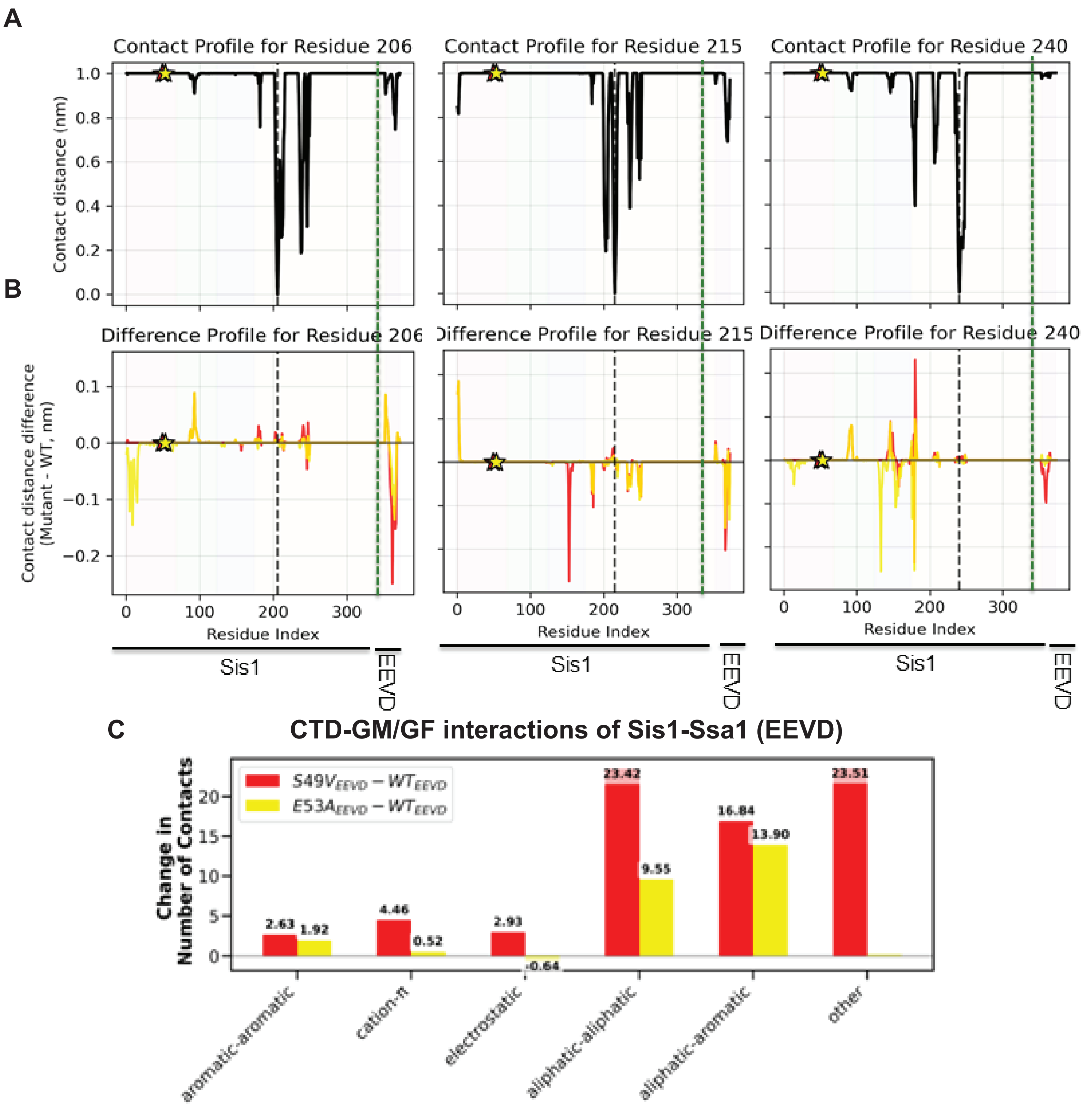


**Supplementary Figure 9. EEVD forms new contacts with Sis1 mutants.** KL divergence analysis (see also Figure 7) and the difference-contact map analysis of Sis1WT-Ssa1 (EEVD) and its mutant complexes showed that the EEVD form new contacts (155, 156, 162, and 168) in mutants. (A) The per-residue contact profile of new contacts in the Sis1WT-EEVD complex shows that these CTD residues bind within the CTD domain. However, they bind weakly to the EEVD motif. The vertical black dashed line represents the position of a specific CTD residue, and the vertical green dashed line represents the position from which EEVD residues start in the complex. (B) Residues 155, 156, 162, and 168 exhibit increased contact persistence with EEVD in mutants (negative change in values). Asterisks highlight mutation-adjacent positions. (C) The average number of inter-domain contacts was calculated in Sis1WT/mutants complexed with Ssa1(EEVD), categorized by residue type. Aliphatic-aliphatic and aliphatic-aromatic interactions dominate the binding. The difference in the average number of contacts revealed that both S49V and E53A display a net increase in hydrophobic interactions between GM and CTD domains.

### Supplementary Figure 10

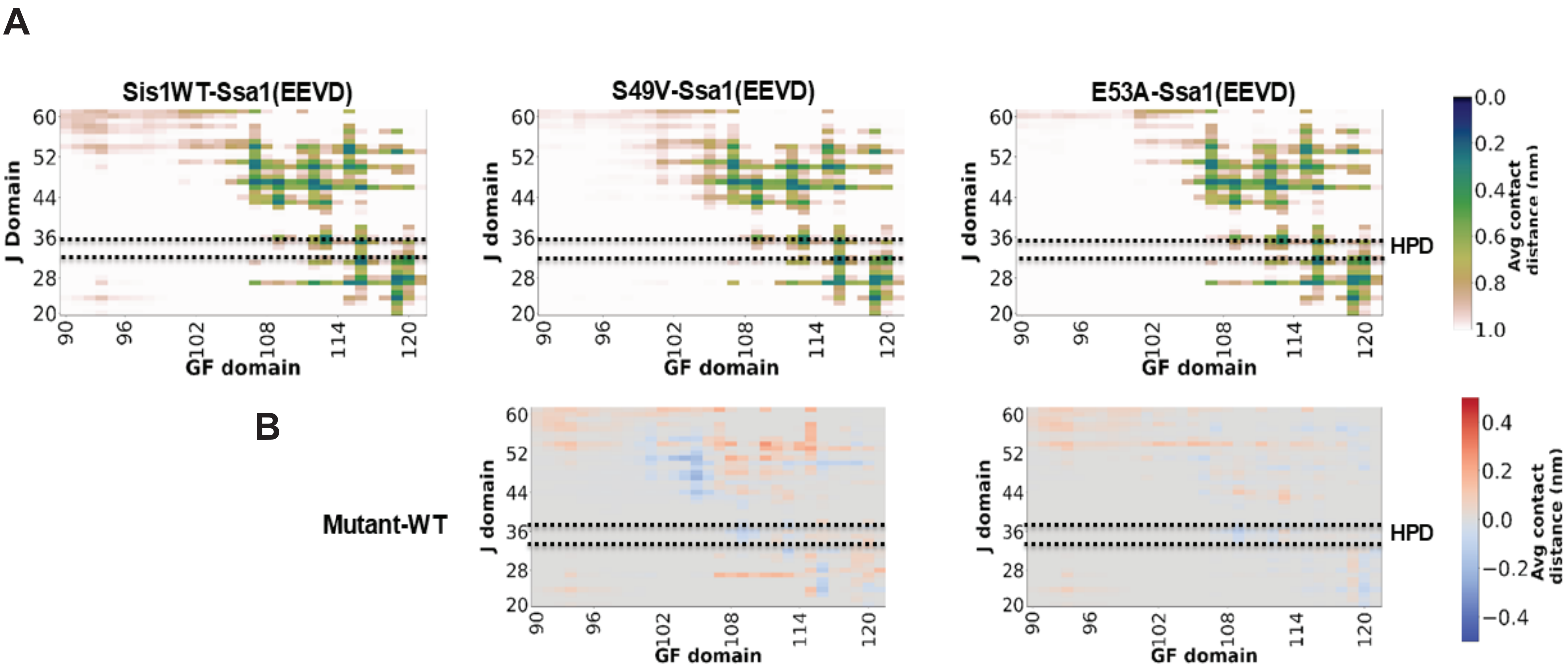


**Supplementary Figure 10. HPD–GF interactions are preserved upon EEVD binding in the absence of the client protein.** (A) Average contact maps of EEVD-bound complexes show stable HPD–GF interactions across WT and mutants. (B) Difference-contact maps confirm no statistically significant alteration in HPD–GF domain interactions due to the EEVD motif alone. line. The dashed line indicates the region of the HPD motif.

#### Supplementary Figure 11

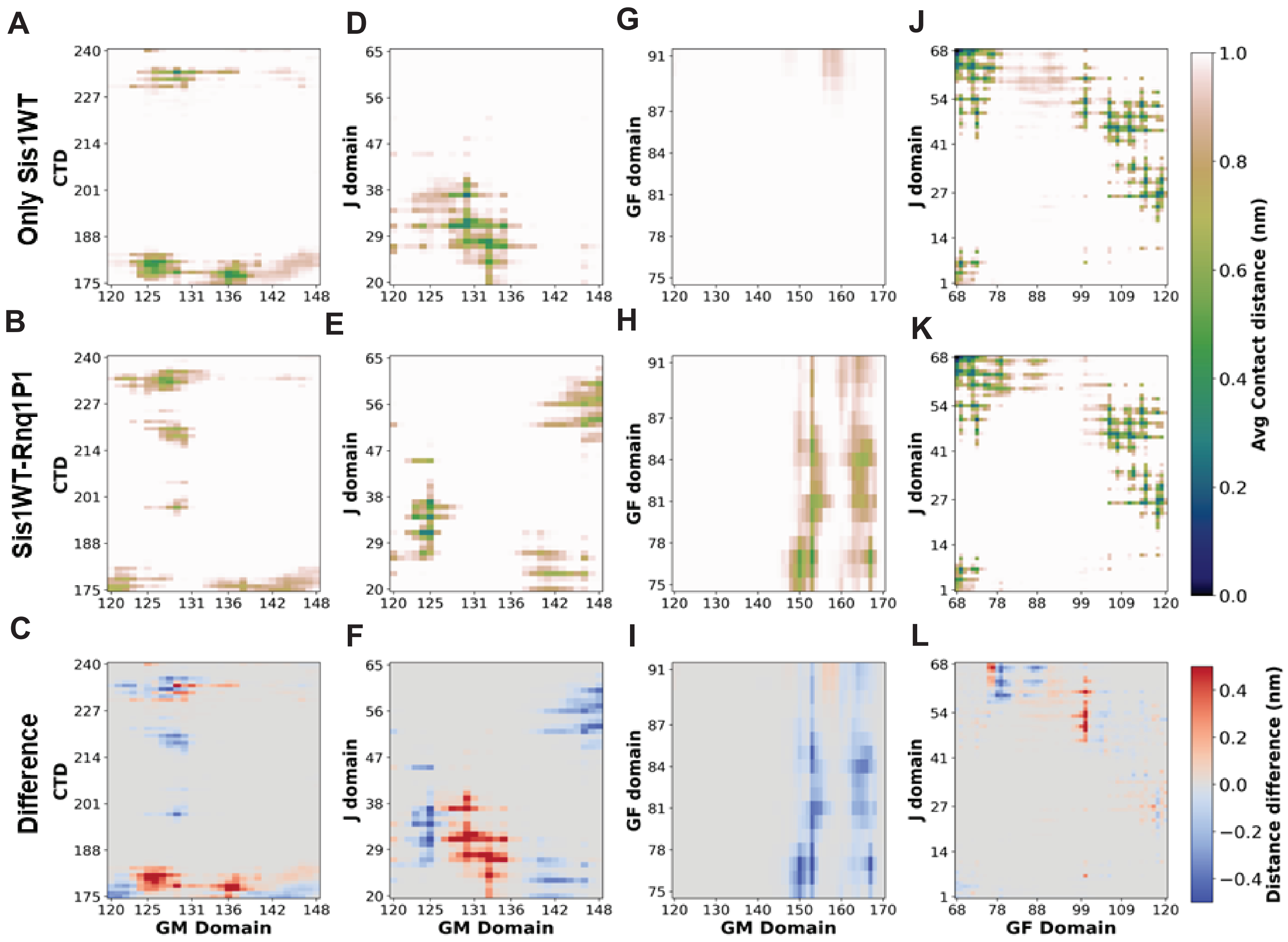


**Supplementary Figure 11. Rnq1 peptide engagement with Sis1WT drives the G-rich region (of the GF domain) to bind J and GM domain.** Zoomed in representation of the average contacts between CTD-GM (A-B); GM-J (D-E); GF-GM (G-H) and GF-J (J-K) in Sis1 WT and Sis1WT-Rnq1P1 complex. (C and F) Difference-contact maps show that upon Rnq1P1 binding, GM residues lose contact with the CTD and J domain. The G-rich region of the GF domain (75-84) forms new contacts with the GM domain (I) and the J domain (L).

### Supplementary Figure 12

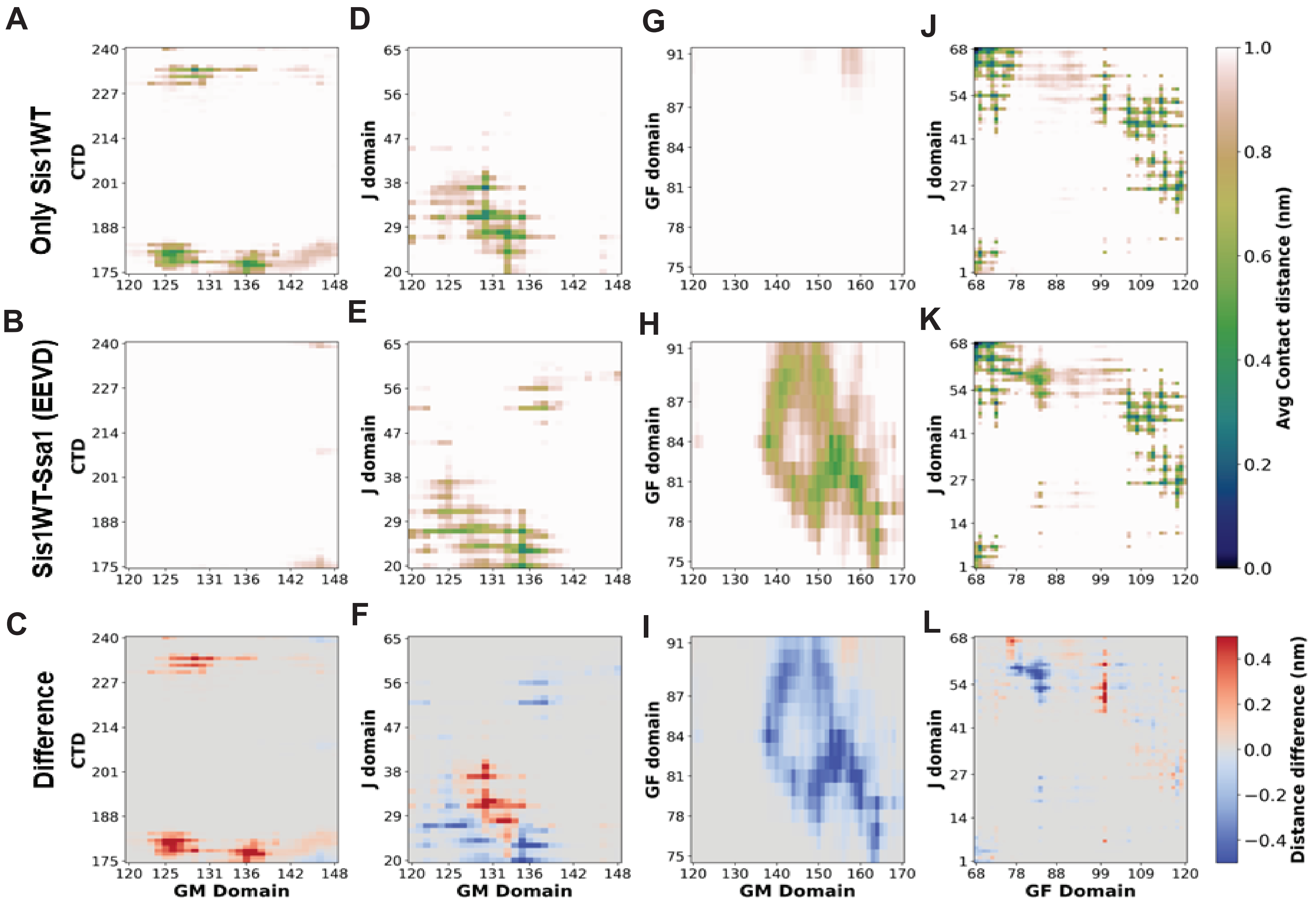


**Supplementary Figure 12 Ssa1 (EEVD) engagement with Sis1WT drives the G-rich region (of the GF domain) to bind J and GM domain.** Zoomed in representation of the average contacts between CTD-GM (A-B); GM-J (D-E); GF-GM (G-H) and GF-J (J-K) in Sis1 WT and Sis1WT-Ssa1(EEVD) complex. (C and F) Difference-contact maps show that upon Ssa1(EEVD) binding, GM residues lose contact with the CTD and J domain. The G-rich region of the GF domain (75-84) forms new contacts with the GM domain (I) and the J domain (L).
